# Scaffold-free human mesenchymal stem cell construct geometry regulates long bone regeneration

**DOI:** 10.1101/785386

**Authors:** Samuel Herberg, Daniel Varghai, Daniel S. Alt, Phuong N. Dang, Honghyun Park, Yuxuan Cheng, Jung-Youn Shin, Anna D. Dikina, Joel D. Boerckel, Marsha W. Rolle, Eben Alsberg

**Author notes:** S.H. and D.V. contributed equally to this work. To whom correspondence should be addressed: Eben Alsberg, Ph.D., Professor; Department of Bioengineering, University of Illinois at Chicago, College of Medicine Research Building, Room 7005, 909 S. Wolcott Avenue, Chicago, IL 60612, USA;.

## Abstract

Scaffold-based bone tissue engineering approaches frequently induce repair processes dissimilar to normal developmental programs. In contrast, biomimetic strategies aim to recapitulate aspects of development through cellular self-organization, morphogenetic pathway activation, and mechanical cues. This may improve regenerative outcome in large long bone defects that cannot heal on their own; however, no study to date has investigated the role of scaffold-free construct geometry, in this case tubes mimicking long bone diaphyses, on bone regeneration. We hypothesized that microparticle-mediated *in situ* presentation of transforming growth factor-β1 (TGF-β1) and bone morphogenetic protein-2 (BMP-2) to engineered human mesenchymal stem cell (hMSC) tubes induces the endochondral cascade, and that TGF-β1 + BMP-2-presenting hMSC tubes facilitate enhanced endochondral healing of critical-sized femoral segmental defects under delayed *in vivo* mechanical loading conditions compared to loosely-packed hMSC sheets. Here, localized morphogen presentation imparted early chondrogenic lineage priming, and stimulated robust endochondral differentiation of hMSC tubes *in vitro*. In an ectopic environment, hMSC tubes formed a cartilage template that was actively remodeled into trabecular bone through endochondral ossification without lengthy predifferentiation. Similarly, hMSC tubes stimulated *in vivo* cartilage and bone formation and more robust healing in femoral defects compared to hMSC sheets. New bone was formed through endochondral ossification in both groups; however, only hMSC tubes induced regenerate tissue partially resembling normal growth plate architecture. Together, this study demonstrates the interaction between mesenchymal cell condensation geometry, bioavailability of multiple morphogens, and defined *in vivo* mechanical environment to recapitulate developmental programs for biomimetic bone tissue engineering.

**Significance Statement:** Engineered bone constructs must be capable of withstanding and adapting to harsh conditions in a defect site upon implantation, and can be designed to facilitate repair processes that resemble normal developmental programs. Self-assembled tubular human mesenchymal stem cell constructs were engineered to resemble the geometry of long bone diaphyses. By mimicking the cellular, biochemical, and mechanical environment of the endochondral ossification process during embryonic development, successful healing of large femoral segmental defects upon implantation was achieved and the extent was construct geometry dependent. Importantly, results were obtained without a supporting scaffold or lengthy predifferentiation of the tubular constructs. This indicates that adult stem/progenitor cells retain features of embryonic mesenchyme, and supports the concept of developmental engineering for bone regeneration approaches.

## Introduction

Long bones of the appendicular skeleton such as the femur provide load bearing structure and carry out critical metabolic functions (1). Mesenchymal cell condensation in the early limb bud initiates long bone morphogenesis to form the cartilage anlage that gives rise to endochondral bone formation (2, 3). This process is dependent on local morphogen gradients and mechanical forces *in utero* (reviewed in (4)). Postnatal bone retains the remarkable ability to heal via endochondral ossification without fibrous scarring (5–7); however, perturbations of the fracture site can interfere with the repair process when bone defects reach a critical size, which can contribute to non-union (8).

Traditional scaffold-based bone tissue engineering approaches aim to create implantable constructs through combination of biocompatible scaffolds, regeneration-competent cells, and/or bioactive cues (9). Owing to the anatomy of the femoral diaphysis, studies have explored preformed tubular constructs in ectopic and orthotopic environments. Natural coral (10–12) and synthetic polymers (13, 14) have been investigated as tubular scaffold materials for mesenchymal stem cells (MSCs). Furthermore, cell-free tubular composites of synthetic polymers and bone morphogenetic protein-2 (BMP-2) or BMP-7 (15–18) have been tested. Limitations of such strategies include (i) the use of scaffolds designed to match the properties of mature rather than developing tissues, (ii) the necessity to pre-culture MSCs for 2-3 weeks in induction medium to stimulate osteogenic pre-differentiation, and (iii) the delivery of a single morphogen.

Biomimetic bone tissue engineering approaches that recapitulate normal tissue development and remodeling have gained considerable interest as of late (19). Scaffold-free cartilage templates derived from self-assembled human MSC (hMSC) condensations have been shown to progress through *in vivo* endochondral ossification, contingent on *in vitro* morphogen priming (20–26). Limitations of this approach include (i) the necessity to pre-culture hMSC condensations for ≥3 weeks in induction medium supplied with morphogens (e.g., transforming growth factor-β1 (TGF-β1)) to stimulate cartilage template formation, and (ii) and the inability to generate tubular tissue constructs. To that end, we have recently reported a modular, scalable, scaffold-free system for engineering hMSC rings that can be assembled and fused into tubular structures (27–29). The incorporation of TGF-β1-presenting gelatin microspheres for *in situ* chondrogenic priming (30) and BMP-2-presenting mineral-coated hydroxyapatite microparticles to induce bony remodeling of the cartilaginous template (31) circumvents the need for lengthy pre-differentiation and may enable early *in vivo* implantation.

In this study, we aimed to recapitulate the cellular, biochemical, and mechanical environment of the endochondral ossification process during early limb development for large long bone defect regeneration. We tested the hypotheses that (i) microparticle-mediated *in situ* presentation of TGF-β1 + BMP-2 to engineered hMSC condensate tubes induces the endochondral cascade *in vitro* and in an ectopic environment *in vivo*, and that (ii) the geometry of TGF-β1 + BMP-2-presenting hMSC tubes, stabilized with custom compliant fixation plates that allow delayed initiation of *in vivo* loading, facilitate enhanced endochondral healing of critical-sized femoral segmental defects compared to loosely-packed hMSC sheets.

## Results

### In vitro maturation of engineered tubular condensations

Endochondral bone formation is initiated by condensation and chondrogenic lineage commitment of mesenchymal cells. Therefore, we seeded a suspension of hMSCs and microparticles to locally present 1) TGF-β1, 2) BMP-2, or 3) TGF-β1 + BMP-2 from within the maturing construct in custom agarose molds to facilitate tissue ring self-assembly by day 2, as described recently (28). Tubular hMSC constructs were then created by placing several ring building blocks in direct contact with each other on horizontal glass tubes for an additional 6 d to facilitate tissue fusion (Fig. S1). We first investigated the effects of *in situ* morphogen presentation on hMSC tube lineage commitment at day 8 of culture. No gross morphological differences between groups were observed at this time point (Fig. 1A). Transcript analysis of key differentiation markers, normalized to controls without growth factors, revealed only minimally elevated mRNA expression of the chondrogenic genes sex determining region Y-box 9 (SOX9), aggrecan (ACAN), and collagen type 2A1 (COL2A1), and the early osteogenic gene alkaline phosphatase (ALP) in BMP-2-loaded hMSC tubes compared to TGF-β1-presenting constructs (Fig. 1B). In contrast, TGF-β1 + BMP-2 dual presentation significantly potentiated SOX9, ACAN, COL2A1, and ALP mRNA levels compared to both TGF-β1- or BMP-2-only constructs (Fig. 1B). No differences in Runt-related transcription factor 2 (RUNX2) and collagen type 1A1 (COL1A1) expression were observed across groups (Fig. 1B). Immunoblot analysis showed SMAD3 phosphorylation in TGF-β1-loaded hMSC tubes; p-SMAD5 was not induced similar to controls without growth factor (Fig. 1C-E). In contrast, BMP-2 presentation induced significant SMAD3 phosphorylation and notable SMAD5 phosphorylation compared to TGF-β1-loaded hMSC tubes (Fig. 1C-E). While TGF-β1 + BMP-2 co-presentation did not further augment SMAD3 phosphorylation vs. BMP-2-loaded hMSC tubes, SMAD5 phosphorylation was significantly potentiated compared to either TGF-β1- or BMP-2-only constructs (Fig. 1C-E). Histologically, engineered hMSC tubes displayed comparable 3-dimensional cellular organization across groups with relatively evenly distributed gelatin microspheres and no substantial glycosaminoglycan (GAG) or mineral deposition. Of note, the relatively proportionally dispersed Alizarin Red-staining observed in all groups was localized to incorporated mineral-coated hydroxyapatite microparticles (Fig. 1F; Fig. S2A-C). Together, these findings suggest that while hMSC tubes across all groups were phenotypically undifferentiated at day 8 of culture, *in situ* presentation of TGF-β1, BMP-2, or TGF-β1 + BMP-2 imparted robust chondrogenic lineage priming in a morphogen-dependent manner.

**Fig. 1.**
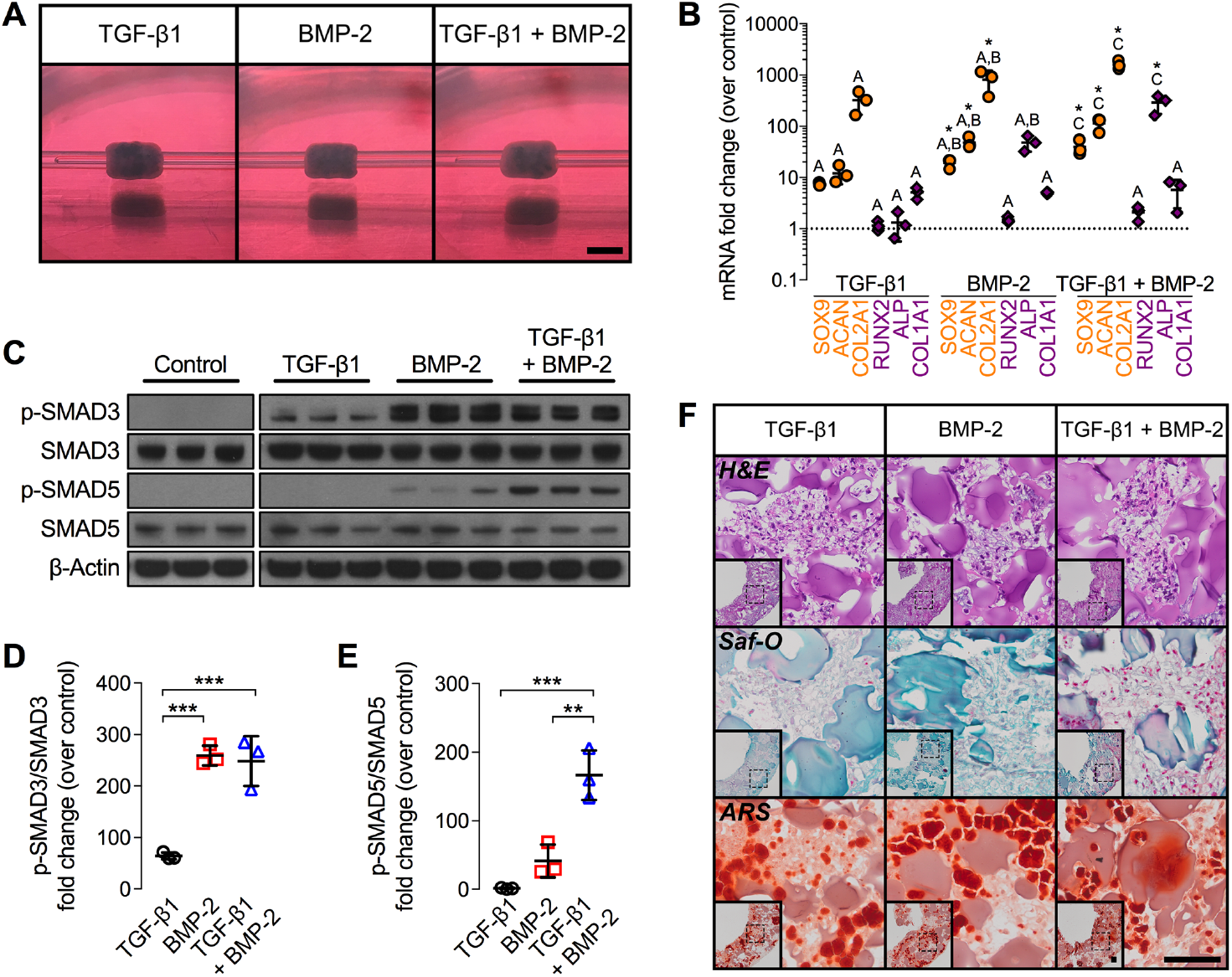
*In vitro* macroscopic, biochemical, molecular, and histological evaluation of engineered hMSC condensate tube early chondrogenic priming. (A) Representative gross macroscopic images of hMSC tubes containing TGF-β1-loaded, BMP-2-loaded, or TGF-β1 + BMP-2-loaded microparticles at day 8. Scale bar, 4 mm. (B) Normalized mRNA fold-change over control of key chondrogenic and osteogenic markers by qRT-PCR (N = 3 per group; *p<0.05 vs. control). (C) Immunoblots and (D) relative fold-change over control of p-SMAD3/SMAD3 and (E) p-SMAD5/SMAD5. β-Actin served as loading control (N = 3 per group; **p<0.01, ***p<0.001). (F) Representative histological Hematoxylin & Eosin (H&E), Safranin-O/Fast green (Saf-O), and Alizarin Red S (ARS) staining of transaxial hMSC tube sections. Scale bars, 100 μm (dotted squares in insets show region of interest in high magnification image). Individual data points shown with mean ± SD. Groups with shared letters have no significant differences. Analyzed by one-way ANOVA with Tukey’s *post hoc* test (p<0.05 or lower considered significant).

Next, we assessed the effects of microparticle-mediated TGF-β1, BMP-2, or TGF-β1 + BMP-2 presentation on stimulating hMSC tube differentiation. Constructs were cultured for 2 weeks in basal medium followed by 3 weeks in osteogenic medium (Fig. S1), conditions previously shown to facilitate robust chondrogenic and osteogenic fate specification in high-density hMSC constructs (28, 32, 33). Gross morphological evaluation revealed comparable width and height of hMSC tubes across groups at 5 weeks (Fig. 2A-C). Histologically, engineered hMSC tubes displayed distinct patterns of tissue differentiation. While all constructs were relatively similar in size and cellularity across groups, presentation of TGF-β1 induced more robust and homogenous GAG deposition, exhibiting cells with chondrocyte-like morphology in Safranin-O-stained regions, compared to BMP-2- or TGF-β1 + BMP-2-loaded constructs (Fig. 2D,E; Fig. S3M,N). Limited mineral deposition was observed with TGF-β1 presentation alone, similar to day 8 constructs. Conversely, BMP-2 and TGF-β1 + BMP-2 presentation induced substantial new mineral deposition by week 5 compared to TGF-β1-only hMSC tubes, consistent with the biochemical assessment (Fig. 2D,E; Fig. S3M,N). Immunohistochemistry revealed robust Col II and Col X staining across groups, indicating the presence of both mature and hypertrophic chondrocytes. In contrast, Col I staining was only observed in BMP-2 and TGF-β1 + BMP-2-loaded hMSC tubes (Fig. S4A-D). Tissue tubes using hMSCs derived from three different donors were prepared for biochemical analysis. The data showed significantly higher GAG/DNA and absolute GAG content with TGF-β1 presentation compared to BMP-2- or TGF-β1 + BMP-2-loaded constructs (Fig. 2F,J; Fig. S3A,E,I). Conversely, BMP-2 and TGF-β1 + BMP-2 presentation promoted significantly higher Ca^2+^/DNA and BMP-2 only presentation promoted significantly greater total Ca^2+^ content relative to TGF-β1-only hMSC tubes (Fig. 2G,K; Fig. S3B,F,J). ALP/DNA and absolute ALP activity was elevated with BMP-2 presentation, reaching significance with dual morphogen delivery vs. TGF-β1 (Fig. 2H,L; Fig. S3C,G,K). No differences in DNA content were noted across groups (Fig. 2I; Fig. S3D,H,L). Together, these findings suggest that *in situ* presentation of TGF-β1, BMP-2, or TGF-β1 + BMP-2 stimulated chondrogenesis, chondrocyte hypertrophy, and osteogenesis indicative of endochondral ossification of hMSC tubes by week 5 *in vitro* in a morphogen-dependent manner.

**Fig. 2.**
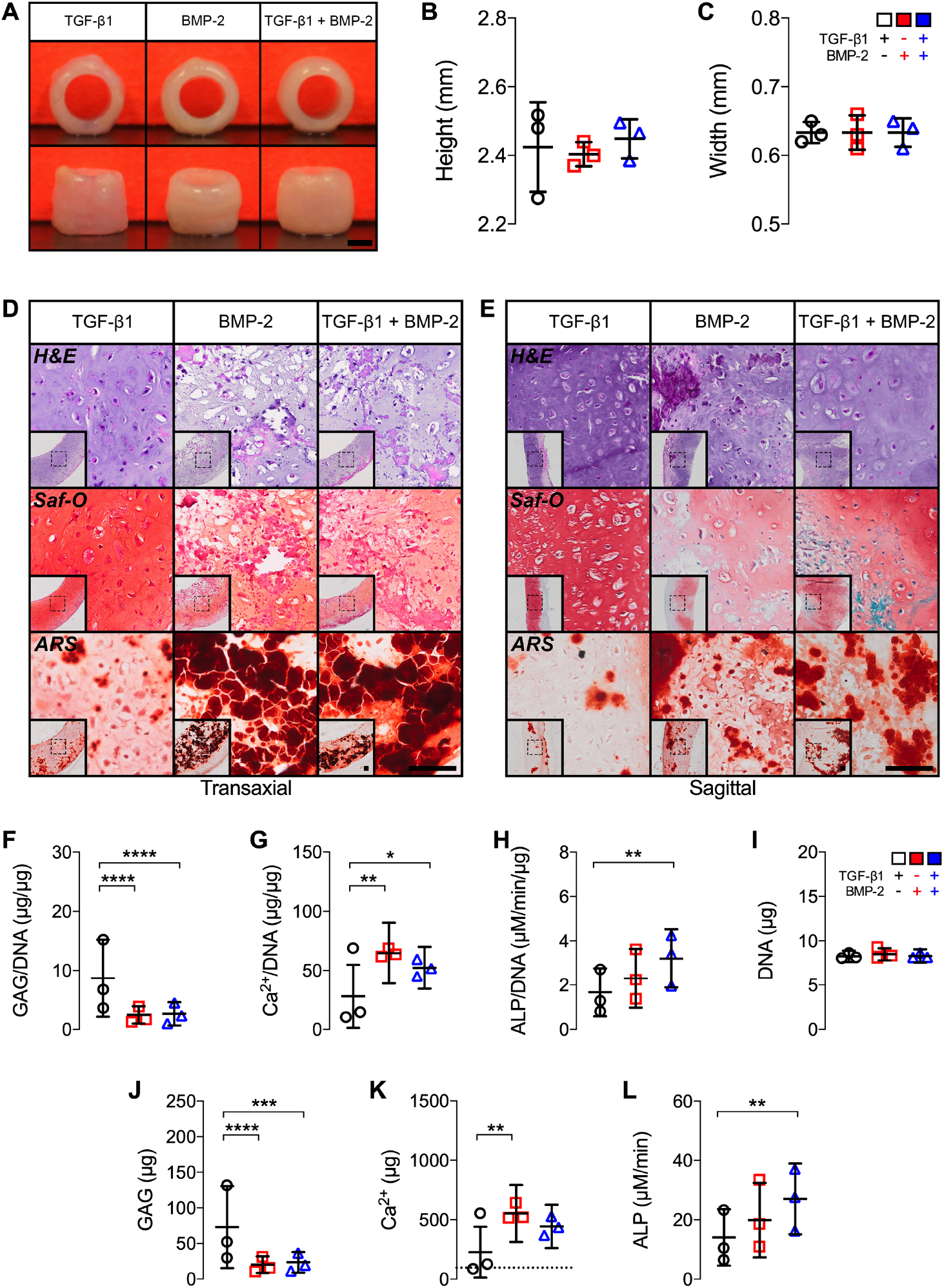
*In vitro* macroscopic, biochemical, and histological evaluation of engineered hMSC condensate tube maturation. (A) Representative gross macroscopic images of hMSC tubes containing TGF-β1-loaded, BMP-2-loaded, or TGF-β1 + BMP-2-loaded microparticles at week 5. Scale bar, 1 mm. (B,C) Quantification of hMSC tube wall width and height (N = 3 per group). (D,E) Representative histological Hematoxylin & Eosin (H&E), Safranin-O/Fast green (Saf-O), and Alizarin Red S (ARS) staining of transaxial and sagittal hMSC tube sections. Scale bars, 100 μm (dotted squares in insets show region of interest in high magnification image). Quantification of (F) GAG/DNA content, (G) Ca^2+^/DNA content, (H) ALP activity/DNA, (I) DNA content, (J) GAG content, (K) Ca^2+^ content; dashed line represents amount of Ca^2+^ initially contributed by mineral-coated hydroxyapatite microparticles assuming 100% incorporation, and (L) ALP activity (N = 3 per group; pooled data (average of averages) from three hMSC donors each with 3-4 replicates; *p<0.05, **p<0.01, ***p<0.001, ****p<0.0001). Individual data points shown with mean ± SD. Analyzed by one-way or two-way ANOVA with Tukey’s *post hoc* test (p<0.05 or lower considered significant).

### In vivo performance of engineered tubular condensations – ectopic implantation

To determine the ability of local TGF-β1, BMP-2, or TGF-β1 + BMP-2 presentation to modulate tissue formation *in vivo*, we subcutaneously implanted day 8 hMSC tubes in 9-week-old NCr-nude mice. Both TGF-β1- and BMP-2-presenting hMSC tube explants were significantly smaller at week 3 compared to TGF-β1 + BMP-2-loaded constructs, which had retained their average implant size; nearly identical patterns were observed at week 6 (Fig. S5A-C). High resolution *ex vivo* microCT analysis revealed limited ossification of TGF-β1- and BMP-2-presenting hMSC tubes at 3 and 6 weeks. Varying degrees of luminal collapse and substantial shrinkage were observed, consistent with the macroscopic evaluation. In contrast, TGF-β1 + BMP-2-loaded constructs underwent robust ectopic ossification at both time points with well-preserved luminal compartments and mineralization patterns guided by the overall implant geometry (Fig. 3A,B*i,ii*; Videos S1,2). Closer inspection of the trabecular architecture in the center of the TGF-β1 + BMP-2-loaded constructs at week 3 revealed relatively tightly packed, thin trabeculae (Fig. 3C*i,i*\*). Four out of six hMSC tubes further exhibited condensed, shell-like mineralized structures on two opposing sides of the constructs’ outermost layer (Fig. 3C*ii,ii*\*). By week 6, central trabeculae appeared markedly thicker (Fig. 3D*i,i*\*). Eight out of eight hMSC tubes exhibited distinct dense outer mineral shells covering a significant portion of the tubes’ outer surface (Fig. 3D*ii,ii*\*). No such evidence was found in any of the TGF-β1- or BMP-2-presenting constructs at either time point. Morphometric analysis revealed markedly increased bone volume in TGF-β1 + BMP-2-loaded hMSC tubes at 3 weeks compared to TGF-β1- and BMP-2-only constructs, which did not differ from each other (Fig. 3E). By week 6, TGF-β1 + BMP-2-loaded tubes exhibited significantly greater bone volume relative to TGF-β1- and BMP-2-only constructs (Fig. 3E). Trabecular number was significantly increased in TGF-β1 + BMP-2-loaded tubes compared to TGF-β1- and BMP-2-only constructs at both time points (Fig. 3F). No differences in trabecular thickness were noted across groups at 3 weeks; however, both BMP-2- and TGF-β1 + BMP-2-loaded tubes exhibited markedly elevated trabecular thickness relative to TGF-β1-only constructs at 6 weeks, reaching significance for BMP-2 vs. TGF-β1 (Fig. 3G). Trabecular separation was markedly decreased in TGF-β1 + BMP-2-loaded tubes compared to TGF-β1- and BMP-2-only constructs at both time points, reaching significance vs. TGF-β1 (Fig. 3H).

**Fig. 3.**
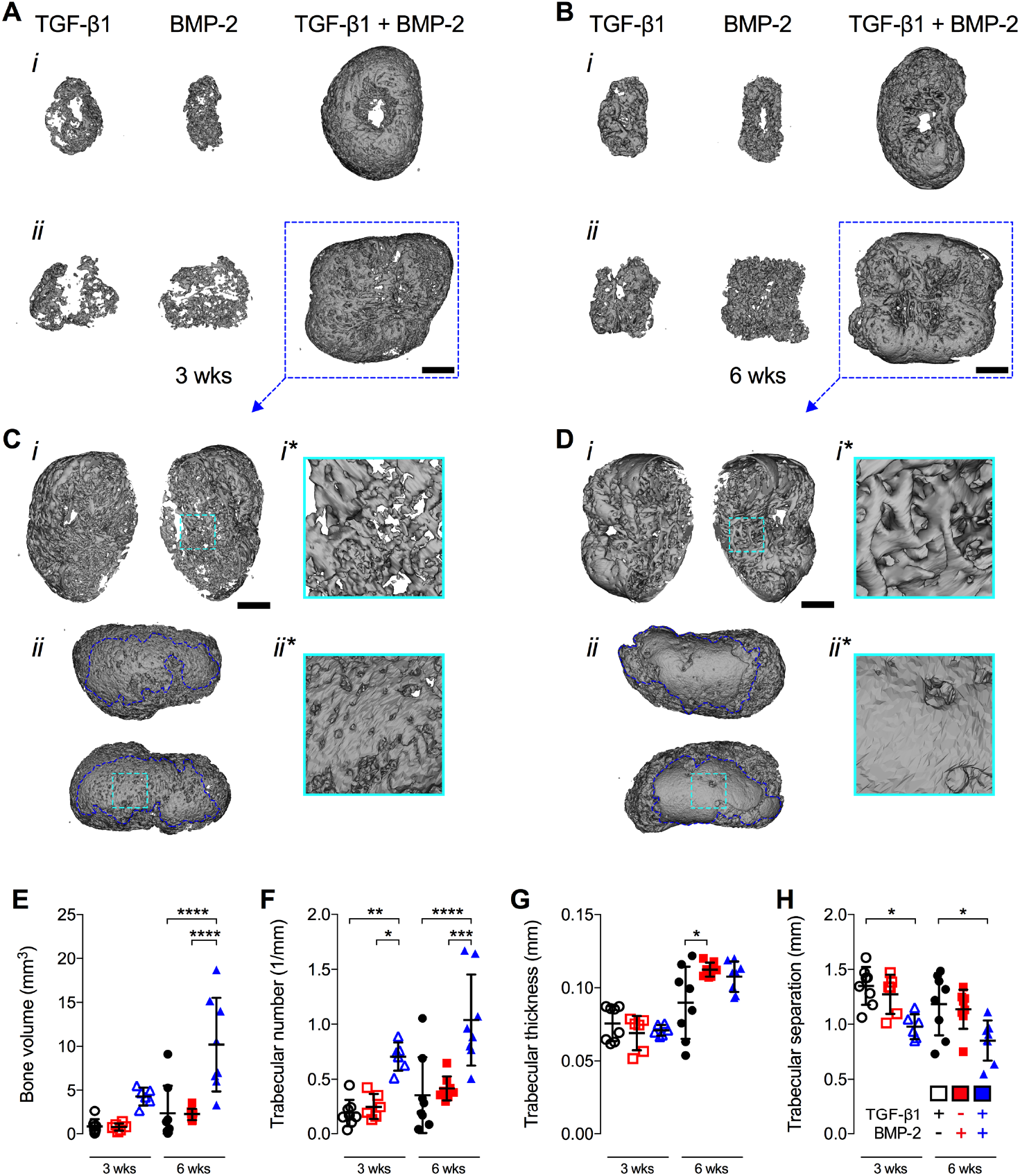
*Ex vivo* microCT evaluation of ectopic bone induced by engineered hMSC condensate tubes. (A,B) Representative 3-D microCT reconstructions of hMSC tube explants containing TGF-β1-loaded, BMP-2-loaded, or TGF-β1 + BMP-2-loaded microparticles at weeks 3 and 6, selected based on mean bone volume. Scale bars, 2 mm. (*i*) top view; (*ii*) side view. (C,D) Multiple angle views and close-ups of (*i,i*\*) trabecular bone and (*ii,ii*\*) cortical bone-like shell architecture in hMSC tube explants containing TGF-β1 + BMP-2-loaded microparticles (dotted blue squares in (A,B)) at weeks 3 and 6. Scale bars, 1 mm. Morphometric analysis of (E) bone volume, (F) trabecular number, (G) trabecular thickness, and (H) trabecular separation at weeks 3 and 6 (N = 6-8 per group). Individual data points shown with mean ± SD. Analyzed by two-way ANOVA with Tukey’s *post hoc* test (p<0.05 or lower considered significant).

Histologically, hMSC tube explants displayed distinct tissue patterns. TGF-β1-loaded constructs induced limited cartilage and bone formation at week 3 *in vivo* (Fig. 4A; Fig. S5D). Presentation of BMP-2 exerted similar effects upon tube implantation; however, cartilage tissue was somewhat less pronounced whereas bone formation was more robust vs. TGF-β1-loaded constructs at week 3 (Fig. 4A; Fig. S5D), consistent with the microCT evaluation. Strikingly, TGF-β1 + BMP-2-loaded hMSC tubes induced significant cartilage and bone formation guided by the implant geometry. Zonal cartilage containing both mature and hypertrophic chondrocytes with prominent GAG matrix were embedded in trabecular bone with osteoid occupying the transitional region, suggestive of endochondral ossification (Fig. 4A; Fig. S5D). Nearly identical tissue formation patterns in a morphogen-dependent manner were observed at week 6 (Fig. 4B; Fig. S5E). Across groups and time points, new tissue was largely comprised of human cells, as evidenced by *in situ* hybridization for human Alu repeats, while a mix of human and mouse cells contributed to the fibrous tissue infiltrating portions of the luminal space as well as surrounding the hMSC tube constructs (Fig. 4C,D). Immunohistochemistry corroborated the histological findings at 3 and 6 weeks. Col II and Col X staining as proxy for identifying mature and hypertrophic chondrocytes, respectively, was most intense in areas with prominent GAG matrix, whereas Col I staining was strongest in regions with extensive mineral deposition (Fig. S6A-D). Together, these findings suggest that localized presentation of TGF-β1, BMP-2, or TGF-β1 + BMP-2 stimulated *in vivo* cartilage and bone formation in hMSC tubes in a morphogen-dependent manner that was guided by the implant architecture. Importantly, TGF-β1 + BMP-2-loaded constructs formed a robust cartilaginous template that was actively remodeled into mineralized trabecular bone tissue through endochondral ossification in an ectopic environment. The presence of cortical bone-like compartments further suggests engagement of intramembranous ossification. Therefore, for subsequent studies we focused on dual morphogen presentation.

**Fig. 4.**
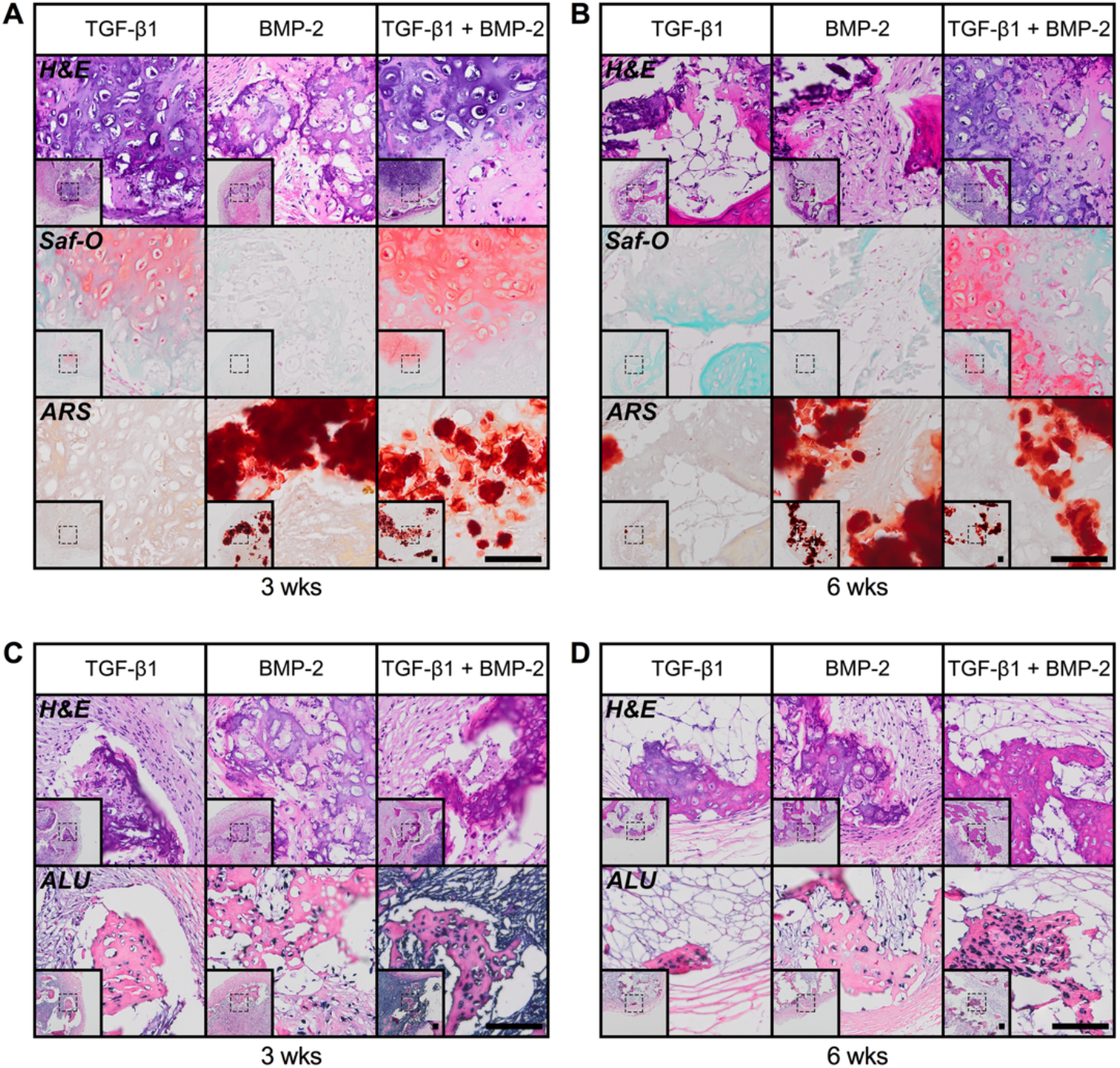
*Ex vivo* histological evaluation of ectopic bone tissue induced by engineered hMSC condensate tubes. (A,B) Representative histological Hematoxylin & Eosin (H&E), Safranin-O/Fast green (Saf-O), and Alizarin Red S (ARS) staining of sagittal sections of hMSC tube explants containing TGF-β1-loaded, BMP-2-loaded, or TGF-β1 + BMP-2-loaded microparticles at weeks 3 and 6. (C,D) Representative H&E staining and *in situ* hybridization for human Alu repeats of transaxial hMSC tube explant sections at weeks 3 and 6. Scale bars, 100 μm (dotted squares in insets show region of interest in high magnification image).

### In vivo performance of engineered tubular condensations – orthotopic implantation

To evaluate the role of engineered condensation geometry on induction of orthotopic bone regeneration, day-8 TGF-β1 + BMP-2-loaded hMSC tubes were implanted in critical-size femoral segmental defects in 12-week-old athymic nude rats. The hMSC tubes’ regenerative potential was compared to that of day-2 TGF-β1 + BMP-2-loaded hMSC sheets comprising matching cell number and morphogen-loaded microparticles. We previously reported successful calvarial defect healing with dual morphogen-loaded hMSC sheets (33). Here, to maintain shape and facilitate vascular infiltration, constructs were placed into perforated, electrospun polycaprolactone (PCL) nanofiber mesh tubes (17, 34, 35) (Fig. S7A). Femora were stabilized with custom fixation plates that permit dynamic tuning of plate compliance *in vivo* (34, 36–39). Similar to our prior studies, plates were initially implanted in a locked configuration (i.e., 0-4 weeks) for stable fixation, and then they were unlocked at week 4 to initiate ambulatory load transfer (i.e., 4-12 weeks) (Fig. S7B), recently shown to enhance the restoration of bone function using growth factor-loaded hMSC sheets (38, 39).

Recognizing the differences in pre-culture times of the TGF-β1 + BMP-2-loaded condensations prior to implantation, 8 days for hMSC tubes and 2 days for hMSC sheets, we evaluated one randomly selected construct per group prior to implantation. Histologically, hMSC tubes appeared denser with noticeable GAG deposition in the outermost tissue layer, but cells did not display mature chondrocyte morphology (Fig. S8A,B). In contrast, hMSC sheets were more loosely packed within the PCL nanofiber mesh tube and showed no evidence of GAG deposition. In both groups, ARS staining was comparable and limited to incorporated mineral-coated hydroxyapatite microparticles. *In situ* hybridization for human Alu repeats confirmed the cells’ identity (Fig. S8A,B). Immunoblot analysis revealed robust SMAD3 and SMAD5 phosphorylation in both groups, with significantly higher levels in hMSC tubes vs. sheets (Fig. S8C-E).

Longitudinal *in vivo* microCT analysis showed increasing bone formation with both TGF-β1 + BMP-2-loaded hMSC tubes and sheets upon implantation in femoral segmental defects over time. Bone volume levels at week 4 were comparable between groups; however, hMSC tubes continued to stimulate bone formation in a near linear fashion upon load initiation at this time point through week 12, reaching significance compared to week 4 (Fig. 5A,D). In contrast, the rate of hMSC sheet-induced bone formation noticeably decreased after week 4 (Fig. 5A,D). Coinciding with plate unlocking, bone volume accumulation rate with hMSC tubes markedly increased between weeks 4 and 8; opposite trends were noted with hMSC sheets. By week 12, bone volume accumulation rates were comparable between groups (Fig. 5B). None of the defects achieved bridging, defined as mineralized tissue fully traversing the defect, at week 4. Fifty percent of defects implanted with TGF-β1 + BMP-2-loaded hMSC tubes were bridged by week 8, whereas hMSC sheets yielded only 33% bridging. By week 12, defect bridging reached 75% with hMSC tubes exhibiting an increasing trend over time, whereas levels remained unchanged with hMSC sheets (Fig. 5C).

**Fig. 5.**
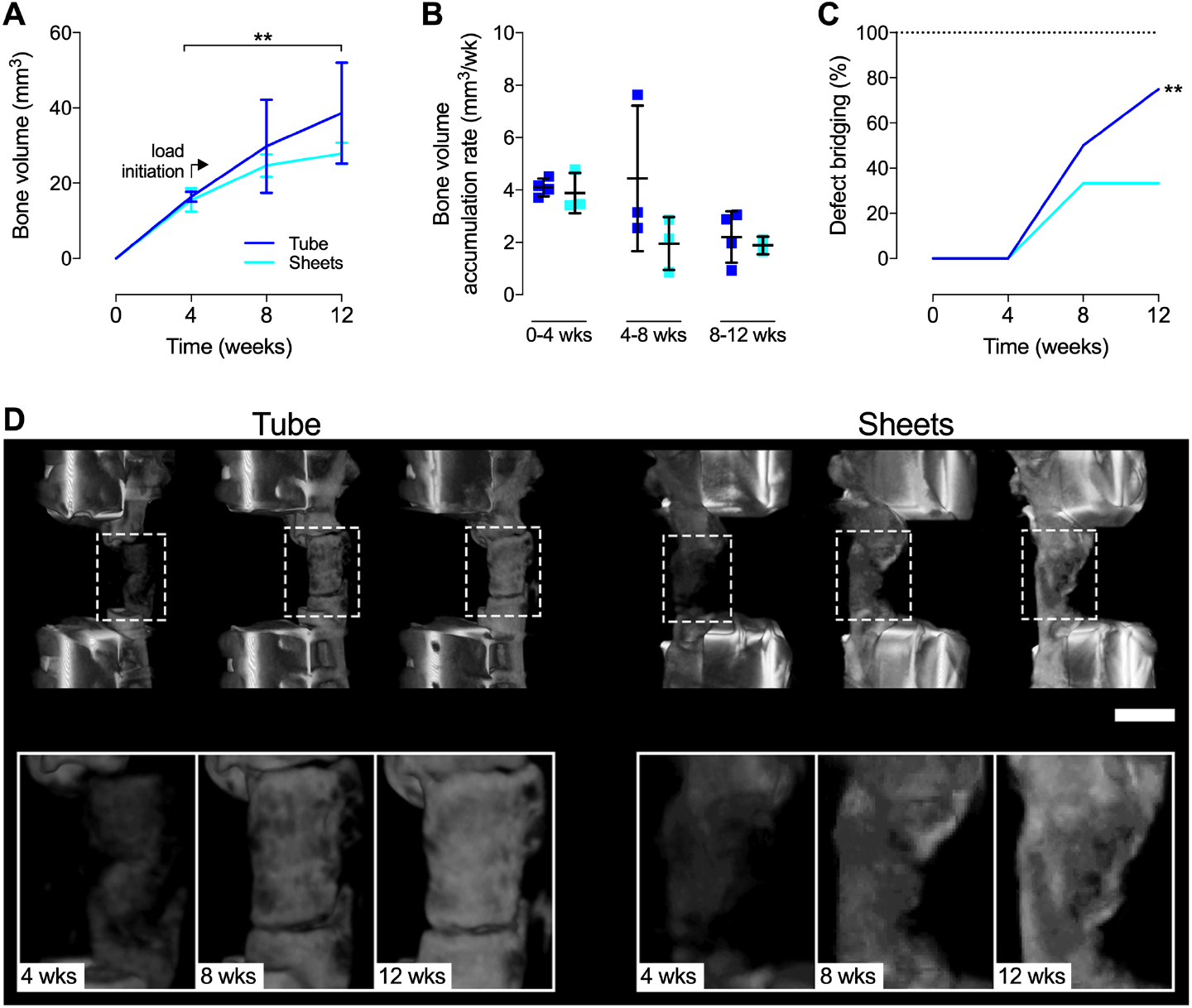
*In vivo* microCT evaluation of femoral defect healing induced by engineered hMSC condensation tubes and sheets. Longitudinal morphometric analysis of (A) bone volume and (B) bone volume accumulation rate at week 4, 8, and 12 in defects implanted with hMSC tubes or sheets containing TGF-β1 + BMP-2-loaded microparticles. Analyzed by two-way ANOVA with Tukey’s *post hoc* test (p<0.05 or lower considered significant). (C) Longitudinal determination of defect bridging, defined as mineral fully traversing the defect (N = 3-4 per group). Significance of trend analyzed by chi-square test (**p<0.01). (D) Representative 3-D microCT defect reconstructions, selected based on mean bone volume. Scale bar, 5 mm. Individual data points shown with mean ± SD.

High-resolution *ex vivo* microCT analysis at 12 weeks demonstrated relatively uniform proximal-to-distal bone distribution in defects treated with either hMSC tubes or sheets with no noteworthy heterotopic ossification (Fig. 6A,B; Video S3). TGF-β1 + BMP-2-loaded hMSC tubes induced greater average bone volume fraction compared to hMSC sheets (Fig. 6C), consistent with the longitudinal *in vivo* assessment. Similarly, while not reaching significance, hMSC tube-treated defects displayed increased average trabecular number and connectivity density vs. hMSC sheets (Fig. 6D; Fig. S9A). The expected inverse relationship was observed for trabecular separation, while trabecular thickness and degree of anisotropy were equivalent across groups (Fig. 6E,F; Fig. S9B). MicroCT-based assessment of structural properties at 12 weeks showed enhanced mean and minimum polar moment of inertia (pMOI) with hMSC tubes compared to hMSC sheets (Fig. S9C,D), theoretically correlating to enhanced torsional strength that was consistent with the bridging data.

**Fig. 6.**
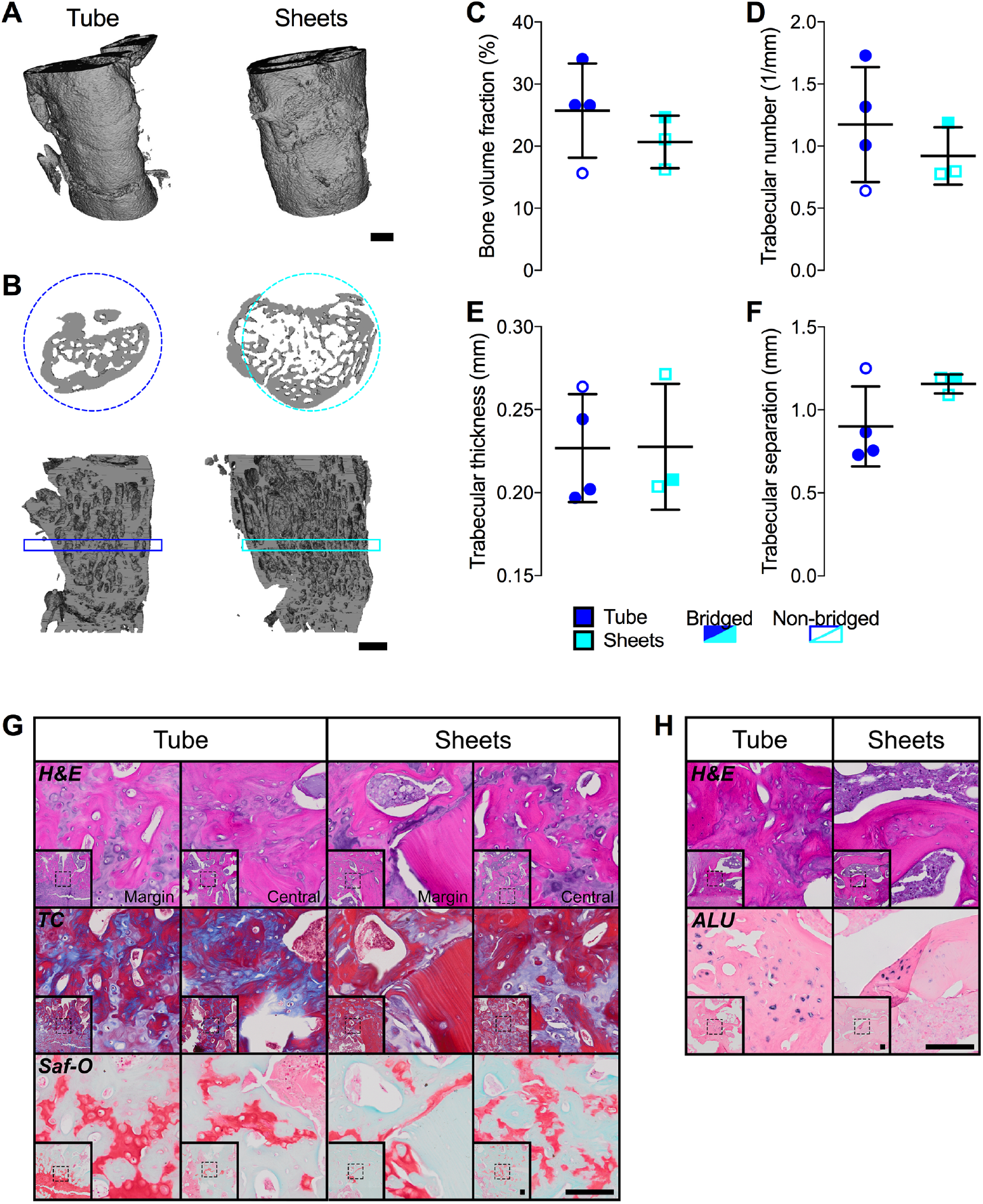
*Ex vivo* microCT and histological evaluation of femoral defect healing induced by engineered hMSC condensate tubes and sheets. (A) Representative 3-D microCT defect reconstructions, implanted with hMSC tubes or sheets containing TGF-β1 + BMP-2-loaded microparticles, with (B) mid-shaft transaxial (top) and sagittal (bottom) sections at week 12, selected based on mean bone volume. Scale bars, 2 mm. Morphometric analysis of (C) bone volume fraction, (D) trabecular number, (E) trabecular thickness, and (F) trabecular separation (N = 3-4 per group). (G) Representative histological Hematoxylin & Eosin (H&E), Masson’s Trichrome (TC), and Safranin-O/Fast green (Saf-O) staining of sagittal defect explant sections showing the defect margin and central defect; images oriented distal-to-proximal from bottom-to-top. (H) Representative H&E staining and *in situ* hybridization for human Alu repeats of sagittal defect explant sections. Scale bars, 100 μm (dotted squares in insets show region of interest in high magnification image). Individual data points shown with mean ± SD. Analyzed by unpaired Student’s t-test (p<0.05 or lower considered significant).

To assess whether *in situ* presentation of TGF-β1 + BMP-2 to engineered hMSC constructs facilitated endochondral regeneration, we performed histological analyses of the defect tissue at week 12. Both hMSC tubes and sheets induced robust ossification; well-defined trabeculae containing lacunae-embedded osteoctyes were found along with cartilaginous structures at different stages of remodeling (Fig. 6G; Fig. S9E). Engineered hMSC tubes promoted formation of growth plate-like, transverse cartilage bands on both the proximal and distal sides connecting the newly formed tissue with the intact bone ends. These remarkable structures exhibited zonal organization of both mature and hypertrophic chondrocytes and were aligned along the principal ambulatory load axis. Prominent GAG matrix was found embedded in trabecular bone with osteoid occupying the transitional region (Fig. 6G; Fig. S9E), indicative of endochondral ossification. No such structures were evident in any of the hMSC sheet-implanted defects. Regenerate bone tissue in both groups at 12 weeks was comprised of rat cells and, as demonstrated by *in situ* hybridization for human Alu repeats, remaining human cells (Fig. 6H). Immunohistochemistry indicated most intense Col II and Col X staining for mature and hypertrophic chondrocytes, respectively, in areas exhibiting robust GAG matrix (Fig. S9F). Together, these findings suggest that localized presentation of TGF-β1 + BMP-2 to hMSC condensations stimulated *in vivo* cartilage and bone formation in an orthotopic environment, independent of implant geometry. However, defined tubular hMSC condensations, exhibiting a greater degree of chondrogenic priming at the time of implantation, facilitated more robust healing compared to loosely-packed hMSC sheets with bony bridging achieved in the majority of critical-size defects. New bone was formed through endochondral ossification in both groups; however, only hMSC tubes induced regenerate tissue that resembled the normal growth plate architecture.

## Discussion

In this study, we report the capacity of scaffold-free hMSC tubes to generate bone tissue *in vivo* through endochondral ossification after only 8 days of pre-culture necessary to allow for tube formation by fusion of individual hMSC rings, and the ability to heal critical-sized rat femoral segmental defects. We employed a biomimetic bone tissue engineering strategy (19) to partially recreate the cellular, biochemical, and mechanical environment of the endochondral ossification process during long bone development. Specifically, we utilized (i) cellular cell self-assembly to form tubular mesenchymal condensations, (ii) microparticle-mediated TGF-β1 + BMP-2 presentation to activate specific morphogenetic pathways *in situ*, and (iii) delayed *in vivo* mechanical loading to augment defect healing. As such, our approach produced a self-organized tubular construct capable of autonomously progressing through formation, maturation, and differentiation into a structured tissue in a manner comparable to normal embryonic development – a concept referred to as developmental engineering (21, 40).

Following a similar biomimetic strategy, others have used scaffold-free, self-assembled hMSC condensations to form cartilage templates that can undergo hypertrophy and progress through endochondral ossification *in vivo* to form ossicles in an ectopic environment, but only after lengthy TGF-β-mediated chondrogenic priming *in vitro* (20–25). The first study that investigated orthotopic implantation of scaffold-free hMSC condensations and showed successful healing of rat femoral segmental bone defects stabilized with internal fixation plates (26) also required lengthy *in vitro* pre-differentiation. We recently demonstrated femoral bone defect healing via endochondral ossification using microparticle-containing hMSC condensate sheets for localized presentation of TGF-β1 (38) or TGF-β1 + BMP-2 (39) that was contingent on *in vivo* loading. No lengthy pre-differentiation of the cellular implants was required. Despite these efforts, *bona fide* tubular tissue constructs have not been developed using such protocols.

This is in contrast to multiple cell-based studies employing traditional bone tissue engineering approaches (9) to form tubular constructs. Natural coral exoskeleton from *Porites* sp. (10–12), and synthetic polymers such as poly lactic-co-glycolic acid (PLGA) (13) or polycaprolactone (PCL) (14) have been investigated as tubular scaffolds for MSCs. Coral/MSC tubes produced mineralized tissue with shape and structure comparable to native long bones *in vivo* in an ectopic environment (11–13), and facilitated near complete healing of sheep metatarsal segmental bone defects stabilized with anchor plates (10). Similarly, implantation of PLGA tubes combined with delayed, minimally-invasive percutaneous injection of MSCs stimulated bone regeneration of sheep tibial segmental defects stabilized with dynamic compression plates (14, 41). While not without merit, these studies were hampered by (i) the use of scaffolds designed to match the properties of mature rather than developing tissues, and (ii) the need for lengthy *in vitro* osteogenic pre-differentiation of MSCs. The risk of poor vascular infiltration of the engineered constructs and subsequent compromised cell survival (42), and differences between the bone repair approaches and normal developmental programs (43) are potential limitations for broad clinical adaptation of such cell-based bone regeneration strategies.

Recently, we reported a modular, scalable, scaffold-free system for engineering hMSC rings that can be assembled and fused into tubular structures (27–29). Here, we also employed gelatin microspheres (30) for early TGF-β1 presentation (32), and mineral-coated hydroxyapatite microparticles (31) for sustained BMP-2 presentation (32), the combination of which exerts potent *in situ* chondrogenic priming effects on early hMSC condensations, consistent with our recent study (28). Importantly, this approach renders time- and cost-intensive pre-differentiation of the cellular constructs obsolete. Both TGF-β1 and BMP-2 play essential roles during embryonic skeletal development and postnatal bone homeostasis; TGF-β signaling is paramount during pre-cartilaginous mesenchymal cell condensation (44), whereas BMP-2 acts upstream of the key transcription factors sex determining region Y-box (SOX) 5, 6, and 9 during early chondrocyte differentiation (45, 46) and is a potent osteogenic growth factor. Here, we showed that localized presentation of TGF-β1 + BMP-2 stimulated cartilage and bone formation in hMSC tubes *in vivo* in an ectopic environment guided by the implant geometry. A robust cartilage template exhibiting zonal architecture with both mature and hypertrophic chondrocytes that was actively remodeled into trabecular bone through endochondral ossification was observed, and the presence of cortical bone-like structures indicated engagement of intramembranous ossification with spatial fidelity. Upon implantation in femoral segmental defects, stabilized with custom internal compliant fixation plates that allow for the delayed commencement of *in vivo* mechanical loading after four weeks of stable fixation (34, 37–39), engineered hMSC tubes induced more robust bone healing compared to loosely-packed hMSC sheets with defect bridging achieved in the majority of critical-size defects. New bone was formed through endochondral ossification in both groups; however, only hMSC tubes induced regenerate tissue that partially resembled the normal growth plate architecture.

Limitations to our study include (i) the difference in pre-culture time between hMSC tubes and sheets, and (ii) the small number of femoral defect samples. To the former, for consistency with previous *in vivo* studies (33, 38, 39), we elected to implant day 2 hMSC sheets in comparison to day 8 hMSC tubes. Condensations were matched in cell number, microparticle concentration, and morphogen dosing; however, the additional six days in culture facilitated greater chondrogenic priming of hMSC tubes vs. sheets. It would be interesting to investigate to what degree this extra pre-culture time contributed to the enhanced regenerative effects in comparison to the implant geometry. To the latter, ten rats required euthanasia 24 h post-surgery; animals presented with marked gastric distention due to ingested hardwood bedding material. This unexpected pica behavior was traced back to buprenorphine administration, as reported by others (47, 48). Consequently, the sample size may have hindered the statistical differences observed between groups in the femoral defect experiment. It would be worthwhile to expand upon the proof-of-principle findings described here in future studies. Furthermore, longer observation times beyond 12 weeks might help address whether tubular condensations are capable of facilitating complete cortical remodeling and re-establishment of a medullary canal.

In conclusion, we describe a biomimetic bone tissue engineering approach that recapitulates certain aspects of the normal endochondral cascade in the developing limb. Implantation of chondrogenically-primed hMSC tubes, achieved through *in situ* morphogen presentation rather than lengthy pre-culture, in large long bone defects that will not heal if left untreated stimulated endochondral ossification for significant bone regeneration with delayed *in vivo* mechanical loading. The tubular hMSC system is of clinical relevance for long bone regeneration, and our findings advance the present understanding in the field of developmental engineering.

## Materials and Methods

Human mesenchymal stromal/stem cells (hMSCs) were obtained from three healthy donors in accordance with the policies of the University Hospitals of Cleveland Institutional Review Board, culture expanded, and used at passage 4. hMSCs were mixed with TGF-β1-loaded gelatin microspheres and BMP-2-loaded mineral-coated hydroxyapatite microparticles, seeded in custom agarose culture wells, and allowed to self-assemble into hMSC rings for 2 days before fusion into tubes by 8 days. hMSC tubes were cultured for 2 weeks in basal medium followed by 3 weeks in osteogenic medium to assess *in vitro* differentiation. All animal experiments were performed in accordance with the National Institutes of Health Guide for the Care and Use of Laboratory Animals, and the policies of the Case Western Reserve University Institutional Animal Care and Use Committee. First, day 8 hMSC tubes were implanted in subcutaneous pouches of 9-week-old nude mice (four implants per mouse) and retrieved after 3 and 6 weeks. Second, day 8 hMSC tubes, contained in polycaprolactone nanofiber mesh tubes, were implanted in critical-sized, bilateral, femoral segmental defects in 12-week-old nude rats (one implant per femur), stabilized with custom compliant fixation plates that allow delayed initiation of *in vivo* loading, and assessed at 4, 8, and 12 weeks. Samples were analyzed by gross macroscopic assessment, micro-computed tomography, biochemical and molecular assays, histology and immunohistochemistry, and *in situ* hybridization. A more detailed description of the methods is included in ***SI Text***.

## Acknowledgments

We thank the staff of the Animal Resource Center at Case Western Reserve University (CWRU) for animal care and husbandry. We also thank the staff of the CWRU Imaging Research Core Facility for *in vivo* imaging support, and Dr. Edward M. Greenfield at the CWRU Orthopaedic Research Facilities for *ex vivo* imaging and biomechanical testing support. We thank Amad Awadallah at the CWRU Histology Core Facility for technical support. We also thank Dr. Xiaohua Yu and Dr. William L. Murphy (University of Wisconsin, Madison, WI) for providing the mineral-coated hydroxyapatite microparticles.

## Author contributions

S.H. and E.A. designed research; S.H., D.V., D.S.A., P.N.D., H.P., Y.C., J-Y.S., and A.D D. performed research; S.H., D.V., and Y.C. analyzed data; and S.H. and E.A. wrote the paper.

