## Supplemental Information for "Scaffold-free human mesenchymal stem cell construct geometry regulates long bone regeneration"

**Supporting Information**

**SI Text**

**Materials and Methods**

***Experimental design.*** The experimental design featured three groups of 3-ring tissue hMSC condensate tubes ($1.2 \times 10^6$ cells) for *in vitro* analyses at 5 weeks: 1) with transforming growth factor-β1 (TGF-β1)-loaded gelatin microspheres (0.4 μg) and unloaded mineral-coated hydroxyapatite microparticles (MCM) **[TGF-β1]**, 2) with unloaded gelatin microspheres (GM) and bone morphogenetic protein-2 (BMP-2)-loaded MCM (1.5 μg) **[BMP-2]**, or 3) with TGF-β1-loaded GM (0.4 μg) and BMP-2-loaded MCM (1.5 μg) **[TGF-β1+BMP-2]** (N = 3-5 per group and donor). Next, 4-ring hMSC condensate tubes ($1.6 \times 10^6$ cells) of the same three groups [1) with TGF-β1-loaded gelatin microspheres (0.5 μg) and unloaded MCM, 2) with unloaded GM and BMP-2-loaded MCM (2.0 μg), or 3) with TGF-β1-loaded GM (0.5 μg) and BMP-2-loaded MCM (2.0 μg)] were implanted subcutaneously to determine ectopic bone formation in male NCr nude mice at 3 and 6 weeks *in vivo* (N = 6-8 per group). Lastly, 8-ring hMSC condensate tubes ($3.2 \times 10^6$ cells) incorporated with TGF-β1-loaded GM (1.0 μg) and BMP-2-loaded MCM (1.0 μg), contained within an electrospun, perforated PCL nanofiber mesh tube, were implanted in femoral segmental defects in male Rowett nude rats to determine longitudinal bone formation over 12 weeks *in vivo* compared to randomly-oriented hMSC sheets at equivalent cell number, microparticle concentration, and growth factor doses (N = 3-4 per group). Limbs were stabilized with axially-compliant fixation plates initially implanted in a locked configuration to prevent loading ($k_{axial} = 250 \pm 35$ N/mm), but after four weeks the plates were surgically unlocked to enable load transfer ($k_{axial} = 8.0 \pm 3.5$ N/mm) (Fig. S7) (1-5).

***hMSC isolation and expansion.*** Human bone marrow-derived mesenchymal stromal/stem cells (hMSCs) were derived from the posterior iliac crest of three healthy donors (M/26, F/25, and M/49 years of age) under a protocol approved by the University Hospitals of Cleveland Institutional Review Board. Cells were isolated using a Percoll (Sigma-Aldrich, St. Louis, MO) density gradient and cultured in low-glucose Dulbecco's modified Eagle's medium (DMEM-LG; Sigma-Aldrich) containing 10% pre-screened fetal bovine serum (FBS; Sigma-Aldrich), and 1% penicillin/streptomycin (P/S; Fisher Scientific). The first media change washed away non-adherent cells, and adherent cells received fresh media supplemented with 10 ng/ml fibroblast growth factor-2 (FGF-2, R&D Systems, Minneapolis, MN) (6, 7).

***Gelatin microsphere synthesis and TGF-β1 loading.*** Gelatin microspheres (GM) (4, 5, 8-11) were synthesized from 11.1% (w/v) gelatin type A (Sigma-Aldrich) using a water-in-oil single emulsion technique and crosslinked for 4 h with 1% (w/v) genipin (Wako USA, Richmond, VA). Hydrated GM were light blue in color and predominantly spherical in shape with an average diameter of 52.9 ± 40.2 μm and a crosslinking density of 25.5 ± 7.0% (11). Growth factor-loaded microspheres were prepared by soaking crosslinked, UV-sterilized GM in a 80 μg/ml solution of rhTGF-β1 (Peprotech, Rocky Hill, NJ) in phosphate-buffered saline (PBS) for 2 h at 37°C. Unloaded GM without growth factor were hydrated similarly using only PBS.

***Hydroxyapatite microparticle mineral coating and BMP-2 loading.*** Mineral-coated hydroxyapatite microparticles (MCM) were kindly provided by Dr. William L. Murphy (University of Wisconsin, Madison, WI). A preparation using low carbonate (4.2 mM $NaHCO_3$) coating buffer and detailed characterization has been reported previously (8, 12). Lyophilized

MCM from the same batch as used in our previous studies (5, 8, 11), were loaded with a 100 µg/ml solution of rhBMP-2 (Dr. Walter Sebald, Department of Developmental Biology, University of Würzburg, Germany; 1.6 or 6.4 µg/mg) in PBS for 4 h at 37°C. BMP-2-loaded MCM were then centrifuged at 800xg for 2 min and washed 2x with PBS. Unloaded MCM without growth factor were incubated with PBS only and treated similarly.

***Preparation of culture wells.*** Custom annular culture well molds (2 mm diameter posts; 3.75 mm wide trough) were 3D printed (Objet260 Connex; Stratasys, Eden Prairie, MN). Polydimethylsiloxane (PDMS; Sylgard 184, Dow Corning, Midland, MI) was cured in the printed molds and served as a negative for casting 2% w/v agarose culture wells (Denville Scientific Inc., Metuchen, NJ). Prior to cell seeding, culture wells were incubated over-night in serum-free, chemically defined basal medium comprised of high-glucose DMEM (Sigma-Aldrich) with 1% $ITS^{+}$ Premix (Corning; Fisher Scientific), 1 mM sodium pyruvate (HyClone; Fisher Scientific), 100 µM non-essential amino acids (Lonza, Basel, Switzerland), 100 nM dexamethasone (MP Biomedicals, Solon, OH), 0.13 mM L-ascorbic acid-2-phosphate (Wako), and 1% P/S (Fisher Scientific) (11, 13, 14).

***Preparation of microparticle-incorporated hMSC condensate tubes.*** Expanded hMSCs ($4.0 \times 10^5$ cells/construct; passage 4) were thoroughly mixed with TGF-β1-loaded GM (0.4 µg/mg; 0.3 mg/construct) and BMP-2-loaded MCM (1.6 or 6.4 µg/mg; 0.08 mg/construct) in basal medium. Fifty microliters of the suspension were seeded in the agarose culture wells as previously described (11, 13, 14) and allowed to self-assemble for 2 days with feeding after 1 day (Fig. S1). Subsequently, tissue rings were removed from the culture wells and transferred to

2 mm diameter borosilicate glass tubes (Adams & Chittenden Scientific Glass, Berkeley, CA) for fusion in 3-, 4-, or 8-ring hMSC condensate tubes. Tissue tubes were cultured horizontally on modified custom polycarbonate holders (11, 13) in 60 mm petri dishes (Corning) containing 10 ml basal medium for 2 weeks, before switching to osteogenic medium for 3 weeks. This time course was previously shown to facilitate robust endochondral ossification (8, 11). Osteogenic medium was identical to basal medium but contained 0.17 mM L-ascorbic acid-2-phosphate and 5 mM β-glycerophosphate (8, 11, 15). Respective media were replaced every 2 d. hMSC tubes incorporated with TGF-β1-loaded GM and unloaded MCM, or unloaded GM and BMP-2-loaded MCM were prepared and cultured in similar fashion. Of note, hMSC condensate tubes incorporated with unloaded GM and unloaded MCM were prepared and served as controls for qRT-PCR and immunoblot analysis. For *in vivo* studies, tubular constructs (ectopic: $1.6x10^6$ cells; orthotopic: $3.2x10^6$ cells) were cultured for a total of 8 days in basal medium prior to implantation, previously shown to be sufficient for fusion into tubes (11).

***Preparation of microparticle-incorporated hMSC condensate sheets.*** Expanded hMSCs ($2.0x10^6$ or $0.6x10^5$ cells/construct; passage 4) were thoroughly mixed with TGF-β1-loaded GM (0.4 μg/mg; 1.5 mg/construct) and BMP-2-loaded MCM (1.6 μg/mg; 0.4 mg/construct) in basal medium. Five hundred or one hundred and fifty microliters of the suspension, respectively, were seeded onto pre-wetted membranes of transwell inserts (3 μm pore size, 12 mm [$2.0x10^6$ cells/construct] or 6.5 mm diameter [$0.6x10^5$ cells/construct]; Corning) and allowed to self-assemble for 2 days (Fig. S7) (5, 16, 17). After 1 day, the medium in the lower compartment was replaced with fresh basal medium and after 2 days, three hMSC condensate sheets (one $2.0x10^6$

cells/construct plus two $0.6 \times 10^5$ cells/construct) were used for implantation for a total of $3.2 \times 10^6$ cells.

***Gross morphological assessment.*** Gross macroscopic images of *in vitro* hMSC condensate tubes at 8 days (N = 3) and subcutaneous hMSC condensate tube explants at 3 and 6 weeks (N = 6-8) were taken with a smartphone camera (Apple Inc., Cupertino, CA) and calibrated using a metric ruler held within the frame of the image. ImageJ software (National Institutes of Health, Bethesda, MD) was used to assess tissue area of the hMSC condensate tubes after initial calibration. Gross macroscopic images of *in vitro* hMSC tubes at 5 weeks (N = 3) were taken with a digital SLR camera (Canon, Melville, NY). Tube wall width and height measurements were obtained using calipers (Fowler High Precision, Newton, MA) at the 12, 4, and 8 o'clock positions (11).

***Quantitative reverse transcription-polymerase chain reaction (qRT-PCR) analysis.*** Day 8 hMSC condensate tubes were homogenized in TRI Reagent (Sigma-Aldrich) for subsequent total RNA extraction and cDNA synthesis (iScript™ kit; Bio-Rad, Hercules, CA). One hundred nanograms of cDNA were amplified in duplicates in each 40-cycle reaction using a Mastercycler (Eppendorf, Hauppauge, NY) with annealing temperature set at 60ºC, SYBR® Premix Ex Taq™ II (Takara Bio Inc., Kusatsu, Shiga, Japan), and custom-designed qRT-PCR primers (Table S1; Life Technologies, Grand Island, NY) (4, 5, 11, 15). Transcript levels were normalized to GAPDH and mRNA fold-change calculated relative to control hMSC condensate tubes without growth factor using the comparative $C_T$ method (18).

**Table S1. Oligonucleotide primer sequences for qRT-PCR.**

| Gene | Sequence (5'-3') | | Accession number |
|---|---|---|---|
| BMPRIA | Fwd | CAGAGATTGGAATCCGCCTGC | NM_004329.2 |
| | Rev | ATCGGGCCGTGCGATCTT | |
| BMPRIB | Fwd | GCAAGCCTGCCATAAGTGAG | NM_001203.2 |
| | Rev | CACAGGCAACCCAGAGTCAT | |
| BMPRII | Fwd | CTGCAAATGGCCAAGCATGT | NM_001204.6 |
| | Rev | ATGGTTGTAGCAGTGCCTCC | |
| TGFBRI | Fwd | ACCCTGCCTAGTGCAAGTTAC | NM_001130916.2 |
| | Rev | AAGCCAAGTTTTCACCCCCA | |
| TGFBRII | Fwd | GTTGGCGAGGAGTTTCCTGTT | NM_001024847.2 |
| | Rev | GTCCTATTACAGCTGGGGCA | |
| SOX9 | Fwd | CACACAGCTCACTCGACCTTG | NM_000346.3 |
| | Rev | TTCGGTTATTTTTAGGATCATCTCG | |
| ACAN | Fwd | TGCGGGTCAACAGTGCCTATC | NM_001135.3 |
| | Rev | CACGATGCCTTTCACCACGAC | |
| COL2A1 | Fwd | GGAAACTTTGCTGCCCAGATG | NM_001844.4 |
| | Rev | TCACCAGGTTCACCAGGATTGC | |
| RUNX2 | Fwd | ACAGAACCACAAGTGCGGTGCAA | NM_001015051.3 |
| | Rev | TGGCTGGTAGTGACCTGCGGA | |
| ALP | Fwd | CCACGTCTTCACATTTGGTG | NM_000478.4 |
| | Rev | GCAGTGAAGGGCTTCTTGTC | |
| COL1A1 | Fwd | GATGGATTCCAGTTCGAGTATG | NM_000088.3 |
| | Rev | GTTTGGGTTGCTTGTCTGTTTG | |
| OSX | Fwd | TGGCTAGGTGGTGGGCAGGG | NM_001173467.2 |
| | Rev | TGGGCAGCTGGGGGTTCAGT | |
| GAPDH | Fwd | GGGGCTGGCATTGCCCTCAA | NM_002046.5 |
| | Rev | GGCTGGTGGTCCAGGGGTCT | |

***Immunoblot analysis.*** Day 8 hMSC condensate tubes (N = 3) were homogenized in CelLytic™ MT lysis buffer (Sigma-Aldrich) supplemented with Halt™ protease and phosphatase inhibitor cocktail (Thermo Scientific). Equal amounts (15 μg) of protein lysates, determined by standard BCA protein assay kit (Pierce; Thermo Fisher Scientific), were subjected to SDS-PAGE using 10% NuPAGE® Bis-Tris gels (Invitrogen; Thermo Fisher Scientific) and transferred to 0.45 μm

PVDF membranes (Millipore, Billerica, MA). Membranes were blocked with 5% bovine serum albumin (BSA) in standard TBST. The phosphorylation of intracellular SMAD3 and SMAD5 was detected using specific primary antibodies (anti-phospho-SMAD3 [ab52903]; anti-phospho-SMAD5 [ab92698]: Abcam, Cambridge, MA) followed by HRP-conjugated secondary antibodies (Jackson ImmunoResearch, West Grove, PA). Subsequently, the blots of phosphorylated SMADs were stripped (Western Blot Stripping Buffer, Pierce; Thermo Fisher Scientific) and re-probed for the detection of the respective total protein (anti-SMAD3 [ab40854]; anti-SMAD5 [ab40771]: Abcam; anti-β-Actin [A1978]: Sigma-Aldrich) with respective HRP-conjugated secondary antibodies (Jackson ImmunoResearch) (4, 5). Bound antibodies were visualized with the ECL detection system (Pierce; Thermo Fisher Scientific) on autoradiography film (Thermo Fisher Scientific). The intensity of immunoreactive bands was quantified using ImageJ software (National Institutes of Health, Bethesda, MD).

***Biochemical analysis.*** Week 5 hMSC condensate tubes (N = 3-4 per donor) were homogenized in papain solution pH 7.4 (Sigma-Aldrich) (19), and assayed for alkaline phosphatase activity (ALP assay kit; Sigma-Aldrich), DNA (PicoGreen kit; Invitrogen, Carlsbad, CA) (20), glycosaminoglycan (GAG) (dimethylmethylene blue kit; Sigma-Aldrich) (21), and calcium content (o-cresophthalein complexone assay kit; Pointe Scientific, Canton, MI) (8, 11).

***Histological and immunohistochemical analysis.*** Week 5 *in vitro* hMSC condensate tubes (N = 3), week 3 and 6 subcutaneous explants (N = 3), and day 8 hMSC condensate tubes and sheets (N = 1) were fixed in 10% neutral buffered formalin (NBF) for 24 h at 4ºC before switching to 70% ethanol. Femora at 12 weeks were fixed in 10% NBF for 72 h at 4ºC and then transferred to

0.25 M ethylenediaminetetraacetic acid (EDTA) pH 7.4 for 14 d at 4ºC under mild agitation on a rocker plate, with changes of the decalcification solution every 3-4 days. Following paraffin processing, 5 μm sections were cut using a microtome (Leica Microsystems Inc., Buffalo Grove, IL) and stained with Hematoxylin & Eosin (H&E), Safranin-O/Fast-green (Saf-O), Alizarin Red S (ARS), or Masson's Trichrome (TC). *In situ* hybridization to detect human Alu repeats was performed using an ISH iVIEW Blue Plus Detection Kit (Ventana Medical Systems Inc., Tucson, AZ) and a BenchMark ULTRA automated IHC/ISH slide staining system (Ventana) according to the manufacturer's instructions. Immunohistochemistry was performed to examine the presence of collagen (Col) types I, II, and X. All sections were deparaffinized and rehydrated with decreasing concentrations of ethanol, and endogenous peroxidase activity was quenched by submerging slides in 30% (v/v) hydrogen peroxide/methanol (1:9) for 10 minutes. For epitope retrieval, sections were digested with pronase (1 mg/ml; Sigma-Aldrich) in PBS at RT for 15 minutes. Rabbit anti-human Col I (ab138492), rabbit anti-human Col II (ab34712), and rabbit anti-human Col X (ab58632; all from Abcam, Cambridge, MA) were used as primary antibodies. Rabbit IgG (Vector Laboratories) served as negative control. The Histostain-Plus Bulk kit (Invitrogen; Thermo Fisher Scientific, Waltham, MA) with aminoethyl carbazole (Invitrogen) was utilized according to the manufacturer's instructions with Fast-green counterstain. Slides were mounted with glycerol vinyl alcohol (Invitrogen), and light microscopy images were captured using an Olympus BX61VS microscope (Olympus, Center Valley, PA) with a Pike F-505 camera (Allied Vision Technologies, Stadtroda, Germany).

***Subcutaneous implantation - ectopic bone formation model.*** Nine-week-old male NCr nude mice were obtained from the Athymic Animal and Xenograft Core Facility at Case Western

Reserve University (CWRU). Animals were anesthetized using 2% isoflurane. Small skin incisions were made on the dorsal side approximately 15 mm from the midline. Subcutaneous pockets were generated using blunt dissection and 4 hMSC condensate tubes (2 per side) were implanted per mouse approximately 30 mm from each other, cranial to caudal. Two constructs from the same group were on the same side in each animal, which were evenly matched with other groups among animals for homogenous pairing distribution between TGF-β1 – BMP-2, TGF-β1 – BMP-2/TGF-β1, and BMP-2 – BMP-2/TGF-β1. The incisions were closed and animals were given subcutaneous injections of 0.1 mg/kg buprenorphine at 0 and 12 h postoperative (post-op). All procedures were performed in strict accordance with the NIH Guide for the Care and Use of Laboratory Animals, and the policies of the CWRU Institutional Animal Care and Use Committee (IACUC) (Protocol No. 2014-0096). Animals were euthanized at 3 and 6 weeks by $CO_2$ asphyxiation followed by cervical dislocation. Skin flaps were opened and hMSC condensate tube explants retrieved.

***Nanofiber mesh production.*** Nanofiber meshes were formed by dissolving 12% (w/v) poly-(ε-caprolactone) (PCL; Sigma-Aldrich) in 90/10 (v/v) hexafluoro-2-propanol/dimethylformamide (Sigma-Aldrich). The solution was electrospun at a rate of 0.75 ml/h onto a static aluminum collector. 9 mm x 20 mm sheets were cut from the product, perforated with a 1 mm biopsy punch (VWR, Radnor, PA) (Fig. S7), and glued into tubes around a 4.5 mm mandrel with UV glue (Dymax, Torrington, CT). Perforated PCL nanofiber mesh tubes were sterilized by 100% ethanol evaporation under UV light over-night and washed 3x with sterile PBS before use. Day 8 microparticle-incorporated hMSC condensate tubes or day 2 hMSC condensate sheets each comprised of $3.2x10^6$ cells were combined into a sterile perforated PCL mesh tube for orthotopic implantation.

***Orthotopic implantation - femoral segmental defect model.*** Critical-sized (8 mm) bilateral segmental defects were created in the femora of 12-week-old male Rowett nude (RNU) rats (Taconic Biosciences Inc., Hudson, NY) using an oscillating saw under isoflurane anesthesia (1-5, 22). Animals received subcutaneous injections of 4 mg/kg lidocaine with 2 mg/kg bupivacaine for local block. Anterolateral incisions were made over the length of each limb, and the *vastus intermedius* and *vastus lateralis* muscles were blunt-dissected to expose the femur. Limbs were stabilized by custom internal fixation plates that allow controlled transfer of ambulatory loads *in vivo* (Fig. S7) (1-5, 22) and secured to the femur by four bi-cortical miniature screws (J.I. Morris Co, Southbridge, MA). Animals were given subcutaneous injections of 0.04 mg/kg buprenorphine every 8 h for the first 24 h post-op with 4 mg/kg carprofen every 24 h for 72 h. In addition, 5 ml of 0.9% NaCl were administered subcutaneously to aid in recovery. All procedures were performed in strict accordance with the NIH Guide for the Care and Use of Laboratory Animals, and the policies of the CWRU IACUC (Protocol No. 2015-0081). Animals were euthanized at 12 weeks by $CO_2$ asphyxiation followed by cervical dislocation. Hind limbs were excised and femora retrieved.

***In vivo microCT.*** *In vivo* microcomputed tomography (microCT) scans were obtained in RNU rats at 4, 8, and 12 weeks to assess longitudinal femoral defect healing according to previously described protocols (4, 5). Data were acquired using an Inveon microPET/CT system (Siemens Medical Solutions, Malvern, PA) at 45 kVp, 0.2 mA, and 35 μm isotropic voxels, and reconstructed using system-default parameters for analyzing bone and accounting for the metal in the fixation plates. DICOM-exported files were processed for 3-D analysis (CTAn software, Skyscan; Bruker, Billerica, MA) using a gauss filter at 1.0 pixel radius and a global threshold range of 28-255 for all samples. This segmentation approach allowed viewing of the normal bone architecture in the binary images as seen in the original reconstructed images (23). One hundred and eighty slices in the center of each defect were analyzed using a 10 mm diameter circle centered on the medullary canal to assess bone volume. A binary bridging score was assigned by two independent, blinded observers, and determined as mineralized tissue fully traversing the defect. Representative 3D images were created using Dornheim Segmenter software (Dornheim Medical Images, Magdeburg, Germany), selected based on average bone volume values.

***Ex vivo microCT.*** *Ex vivo* microCT scans were obtained at 3 and 6 weeks (mouse; ectopic model) or at 4, 8, and 12 weeks (rat; orthotopic model). Data were acquired using a Skyscan 1172 microCT scanner (Skyscan; Bruker) with a 0.5 mm aluminum filter at 75 kVp and 0.1 mA. Ectopic explants wrapped in gauze were placed in a plastic sample holder and scanned in PBS at 10 μm isotropic voxels, 1170 ms integration time, rotation step of 0.5°, and frame averaging of 5. All ectopic explants were scanned within the same container using the same scanning parameters. Femora wrapped in gauze were placed in a plastic sample holder with the long axis oriented parallel to the image plane and scanned in PBS at 20 μm isotropic voxels, 330 ms integration time, rotation step of 0.5°, and frame averaging of 5. All femora were scanned within the same container using the same scanning parameters. All scans were then reconstructed using NRecon software (Skyscan; Bruker) with the same reconstruction parameters (ring artifact reduction of 5, beam hardening correction of 20%). For 3-D analysis (CTAn software, Skyscan; Bruker), a gauss filter at 1.0 pixel radius and a global threshold range of 65-255 were used for all samples. This segmentation approach allowed viewing of the normal bone architecture in the binary images as seen in the original reconstructed images (23). For ectopic explants, varying numbers of slices were analyzed in an 8 mm diameter circle to account for the differences in size. For femora, 325 slices in the center of each defect were analyzed using a 10 mm (total) or 5 mm (defect) diameter circle centered on the medullary canal. Bone volume, bone volume fraction, polar moment of inertia, and the morphometric parameters trabecular number, trabecular thickness, trabecular separation, degree of anisotropy, and connectivity density were calculated (23). Representative 3D images were created using CTAn software (Skyscan), selected based on average bone volume values.

***Statistical analysis.*** Fold-changes over control in mRNA expression and ratios of phosphorylated SMADs/total SMADs, and differences in gross morphological hMSC condensate tube wall width and height, and biochemical markers were analyzed by one-way or two-way analysis of variance (ANOVA) with Tukey's multiple comparison *post hoc* test. Differences in gross morphological tissue area, and *ex vivo* microCT bone volume and 3-D morphometry in hMSC condensate tube explants were analyzed by two-way ANOVA with Tukey's multiple comparison *post hoc* test. Bridging of femoral defects was determined by chi-square test for trend for each group. Differences in longitudinal *in vivo* microCT bone volume and bone volume accumulation rate at 4, 8, and 12 weeks, and *ex vivo* microCT bone volume fraction and 3-D morphometry in femoral defects at 12 weeks were assessed by two-way ANOVA with Tukey's multiple comparison *post hoc* test. All data are shown with mean ± SD, some with individual data points. The significance level was set at $p<0.05$ or lower. Groups with shared letters have no significant differences. GraphPad Prism software v6.0 (GraphPad Software, La Jolla, CA) was used for all analyses.

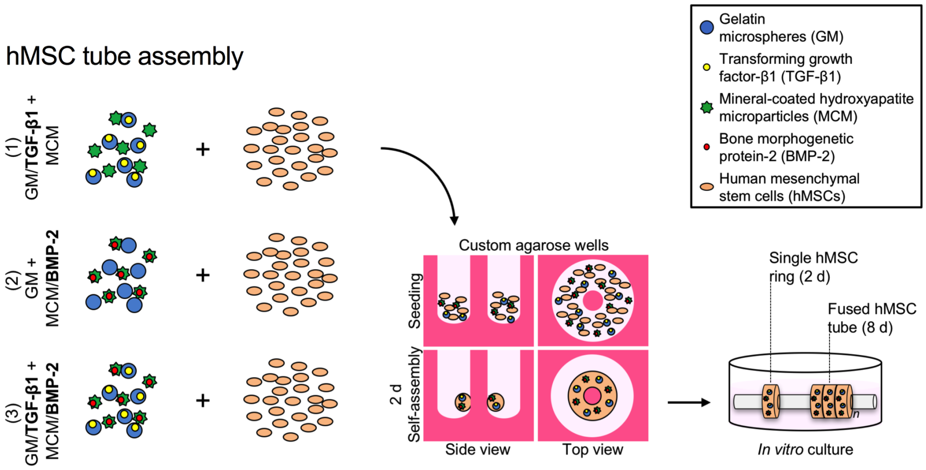


**Fig. S1. Schematic of hMSC condensate tube assembly and culture.** hMSCs were mixed with (1) TGF-β1-loaded gelatin microspheres and unloaded mineral-coated hydroxyapatite microparticles [TGF-β1], (2) unloaded gelatin microspheres and BMP-2-loaded mineral-coated hydroxyapatite microparticles [BMP-2], or (3) TGF-β1-loaded gelatin microspheres and BMP-2-loaded mineral-coated hydroxyapatite microparticles [TGF-β1 + BMP-2], seeded in custom agarose culture wells, and allowed to self-assemble into hMSC rings for 2 days before fusion into tubes by 8 days. hMSC tubes were cultured horizontally on glass tubes for 2 weeks in basal medium followed by 3 weeks in osteogenic medium.

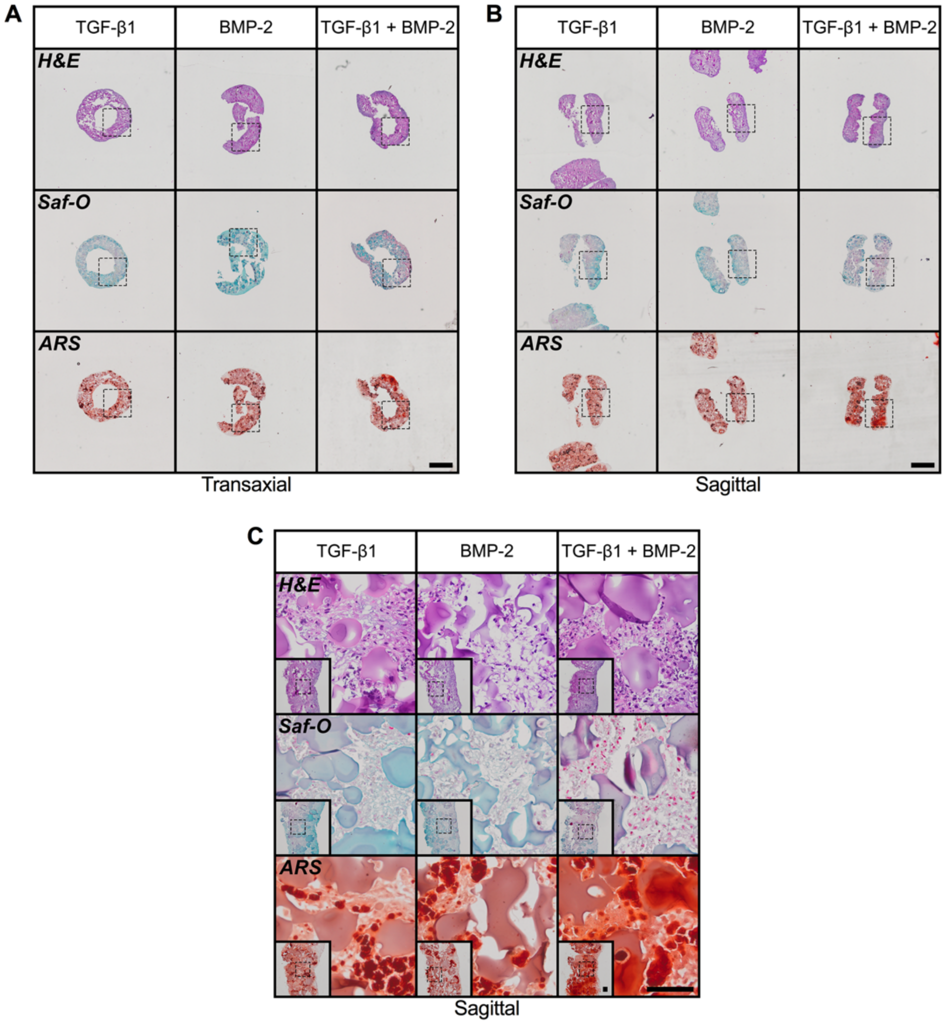


**Fig. S2.** ***In vitro*** **histological evaluation of engineered hMSC condensate tube early chondrogenic priming.** (A,B) Representative histological Hematoxylin & Eosin (H&E), Safranin-O/Fast green (Saf-O), and Alizarin Red S (ARS) staining of transaxial and sagittal sections of hMSC tubes containing TGF-β1-loaded, BMP-2-loaded, or TGF-β1 + BMP-2-loaded microparticles at day 8. Scale bars, 1 mm (dotted squares show areas used in 10x images in Fig. 1F or S2C). (C) Representative H&E, Saf-O, and ARS staining of sagittal hMSC tube sections. Scale bars, 100 µm (dotted squares in insets show region of interest in high magnification image).

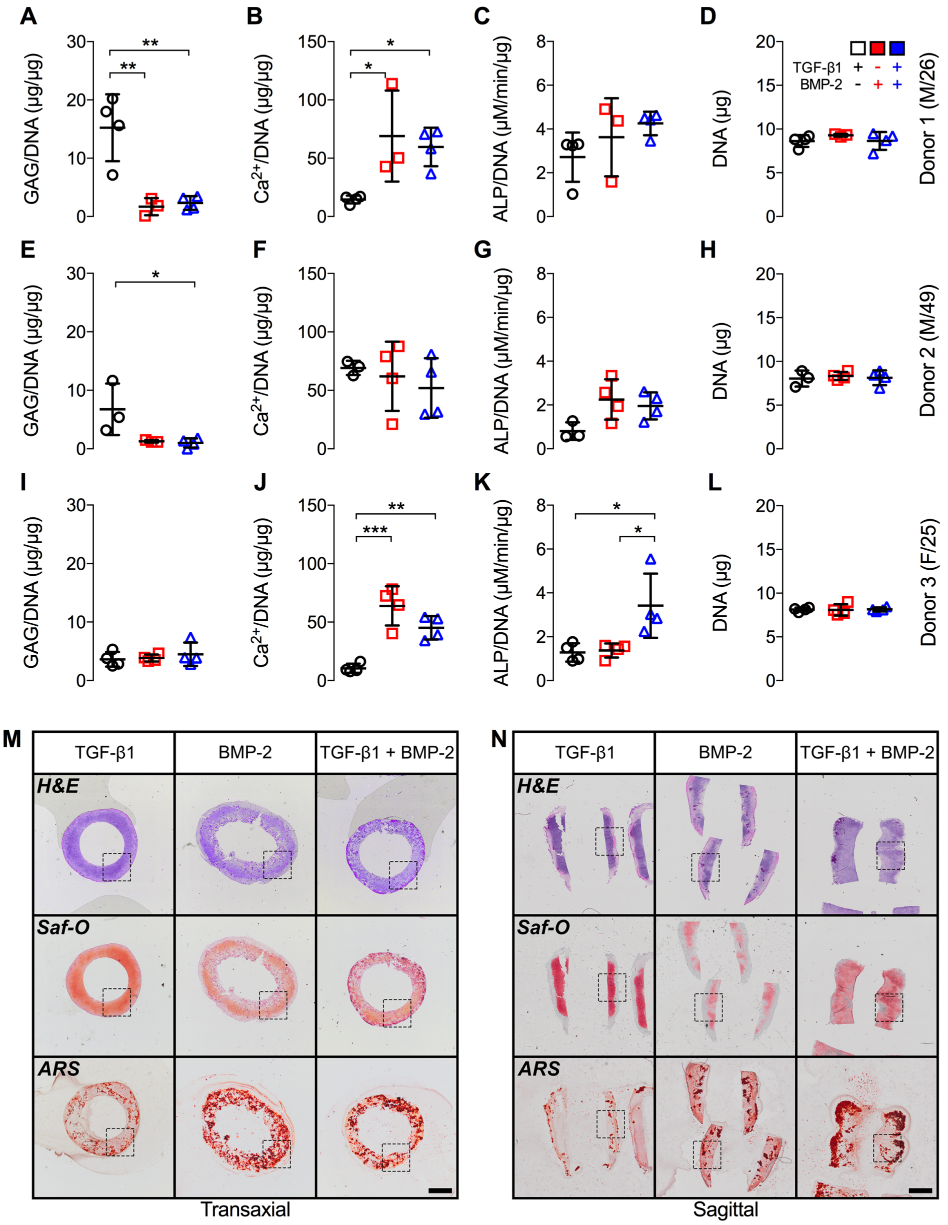


**Fig. S3.** ***In vitro*** **biochemical and histological evaluation of engineered hMSC condensate tube maturation.** Quantification of (A,E,I) GAG/DNA content, (B,F,J) $Ca^{2+}$/DNA content, (C,G,K) ALP activity/DNA, and (D,H,L) DNA content in hMSC tubes, from three separate donors, containing TGF-β1-loaded, BMP-2-loaded, or TGF-β1 + BMP-2-loaded microparticles at week 5 (N = 3-4 per group; *$p<0.05$, **$p<0.01$, ***$p<0.001$). (M,N) Representative histological Hematoxylin & Eosin (H&E), Safranin-O/Fast green (Saf-O), and Alizarin Red S (ARS) staining of transaxial and sagittal hMSC tube sections. Scale bars, 1 mm (dotted squares show areas used in 10x images in Fig. 2K,L). Individual data points shown with mean ± SD. Analyzed by one-way ANOVA with Tukey's *post hoc* test ($p<0.05$ or lower considered significant).

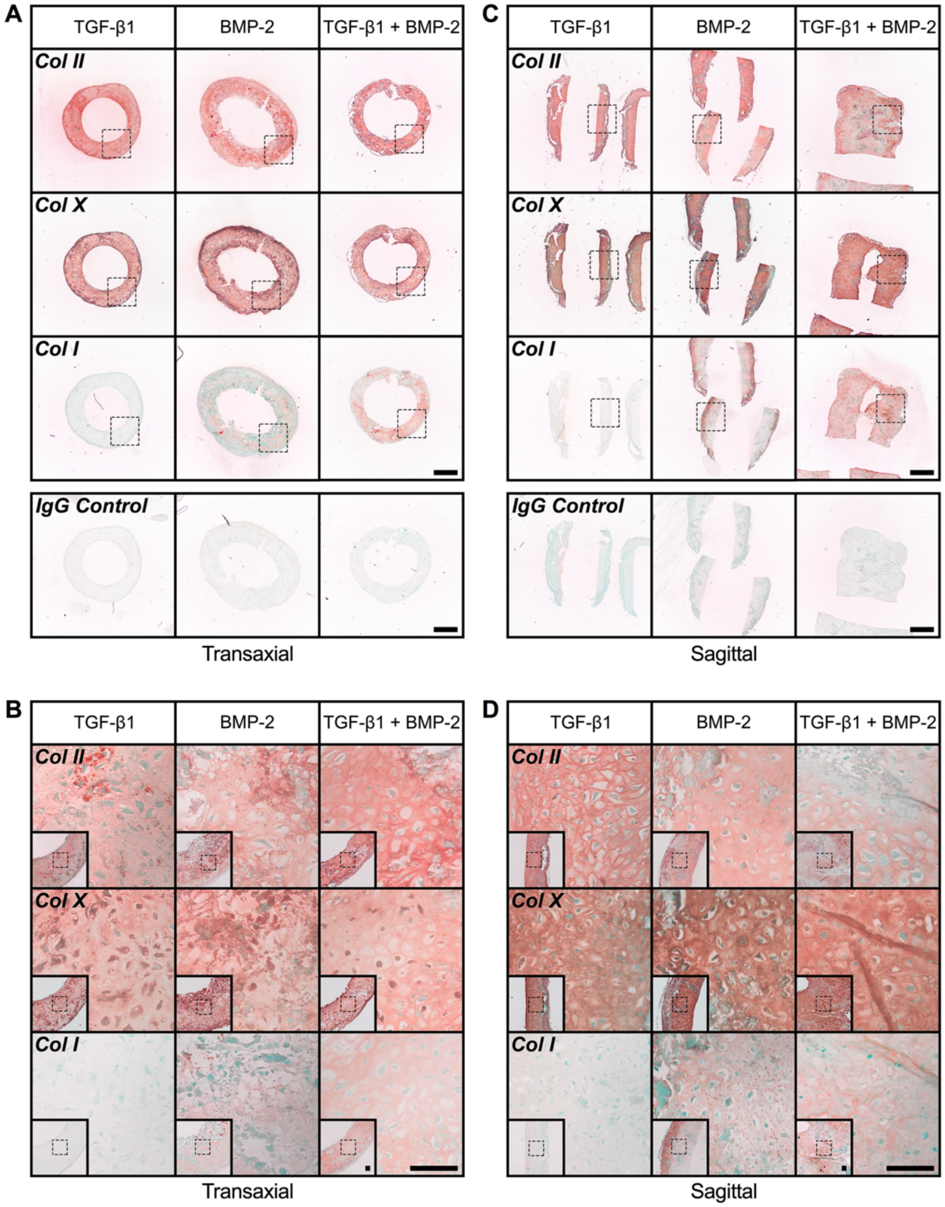


**Fig. S4.** ***In vitro*** **immunohistochemical evaluation of engineered hMSC condensate tube maturation.** (A,C) Representative immunohistochemical collagen (Col) II, Col X, and Col I staining of transaxial and sagittal sections of hMSC tubes containing TGF-β1-loaded, BMP-2-loaded, or TGF-β1 + BMP-2-loaded microparticles at week 5, with representative IgG negative controls. Scale bars, 1 mm (dotted squares show areas used in 10x images in Fig. S4B,D). (B,D) Representative immunohistochemical Col II, Col X, and Col I staining of transaxial and sagittal hMSC tube sections. Scale bars, 100 µm (dotted squares in insets show region of interest in high magnification image).

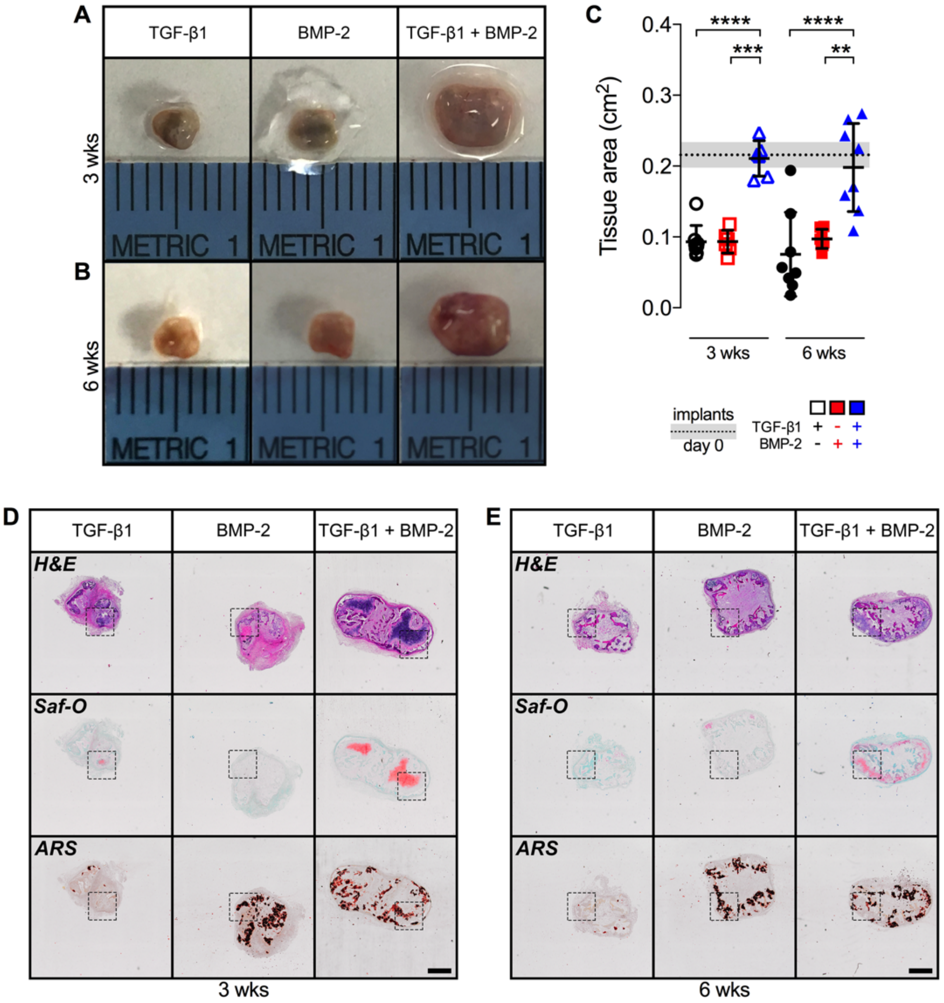


**Fig. S5. *Ex vivo* macroscopic and histological evaluation of ectopic bone tissue induced by engineered hMSC condensate tubes.** (A,B) Representative gross macroscopic images of hMSC tube explants containing TGF-β1-loaded, BMP-2-loaded, or TGF-β1 + BMP-2-loaded microparticles at week 3 and 6, selected based on mean tissue area. (C) hMSC explant tissue area quantification, shown with mean ± SD (gray shading) construct size across groups at the time of implantation (N = 6-8 per group; $^{**}p<0.01$, $^{***}p<0.001$, $^{****}p<0.0001$). (D,E) Representative histological Hematoxylin & Eosin (H&E), Safranin-O/Fast green (Saf-O), and Alizarin Red S (ARS) staining of sagittal hMSC tube explant sections at week 3 and 6. Scale bars, 1 mm (dotted squares show areas used in 10x images in Fig. 4A,B). Individual data points shown with mean ± SD. Analyzed by two-way ANOVA with Tukey's *post hoc* test ($p<0.05$ or lower considered significant).

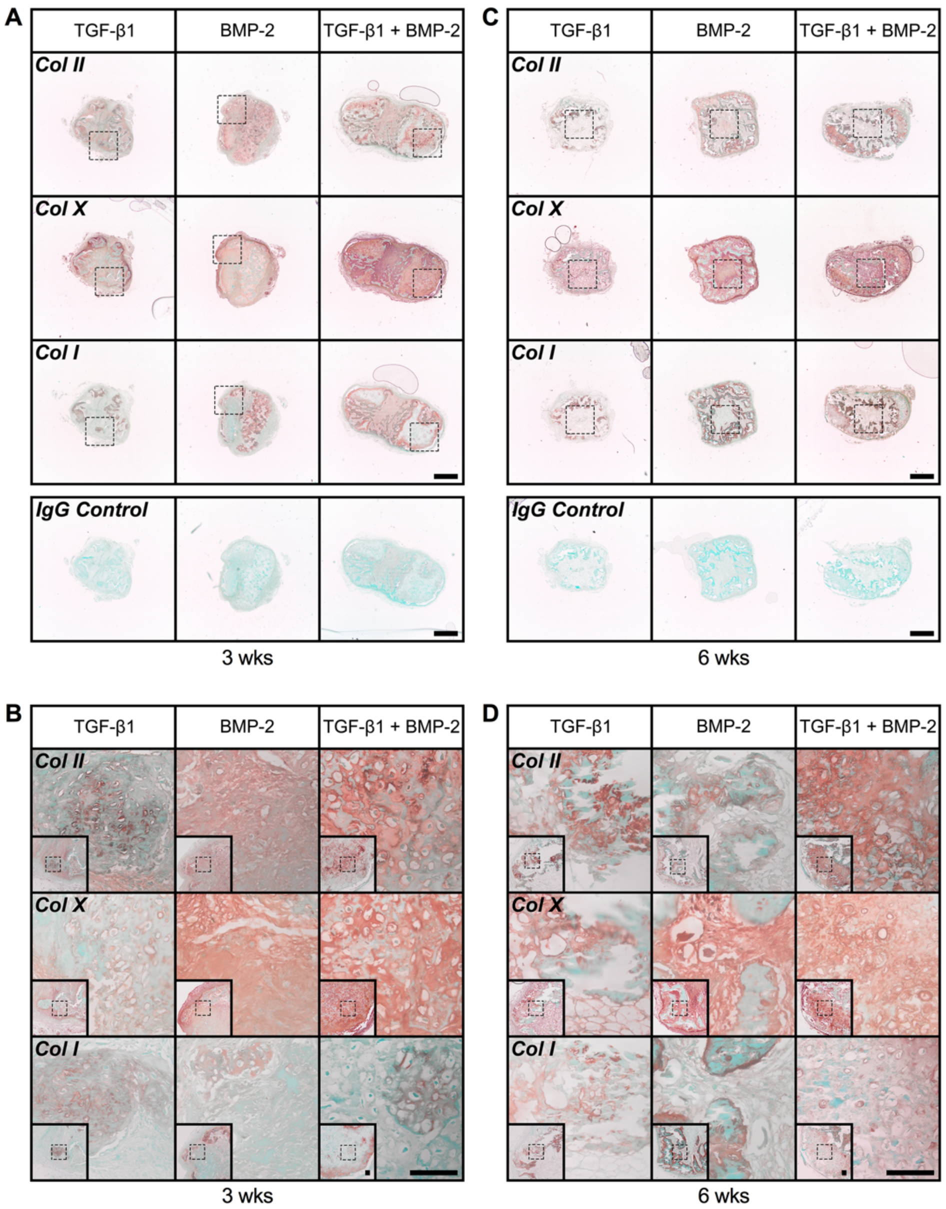


**Fig. S6.** ***Ex vivo*** **immunohistochemical evaluation of ectopic bone tissue induced by engineered hMSC condensate tubes.** (A,C) Representative immunohistochemical collagen (Col) II, Col X, and Col I staining of transaxial sections of hMSC tube explants containing TGF-β1-loaded, BMP-2-loaded, or TGF-β1 + BMP-2-loaded microparticles at week 3 and 6, with representative IgG negative controls. Scale bars, 1 mm (dotted squares show areas used in 10x images in Fig. S6B,D). (B,D) Representative immunohistochemical Col II, Col X, and Col I staining of sagittal hMSC tube explant sections at week 3 and 6. Scale bars, 100 μm (dotted squares in insets show region of interest in high magnification image).

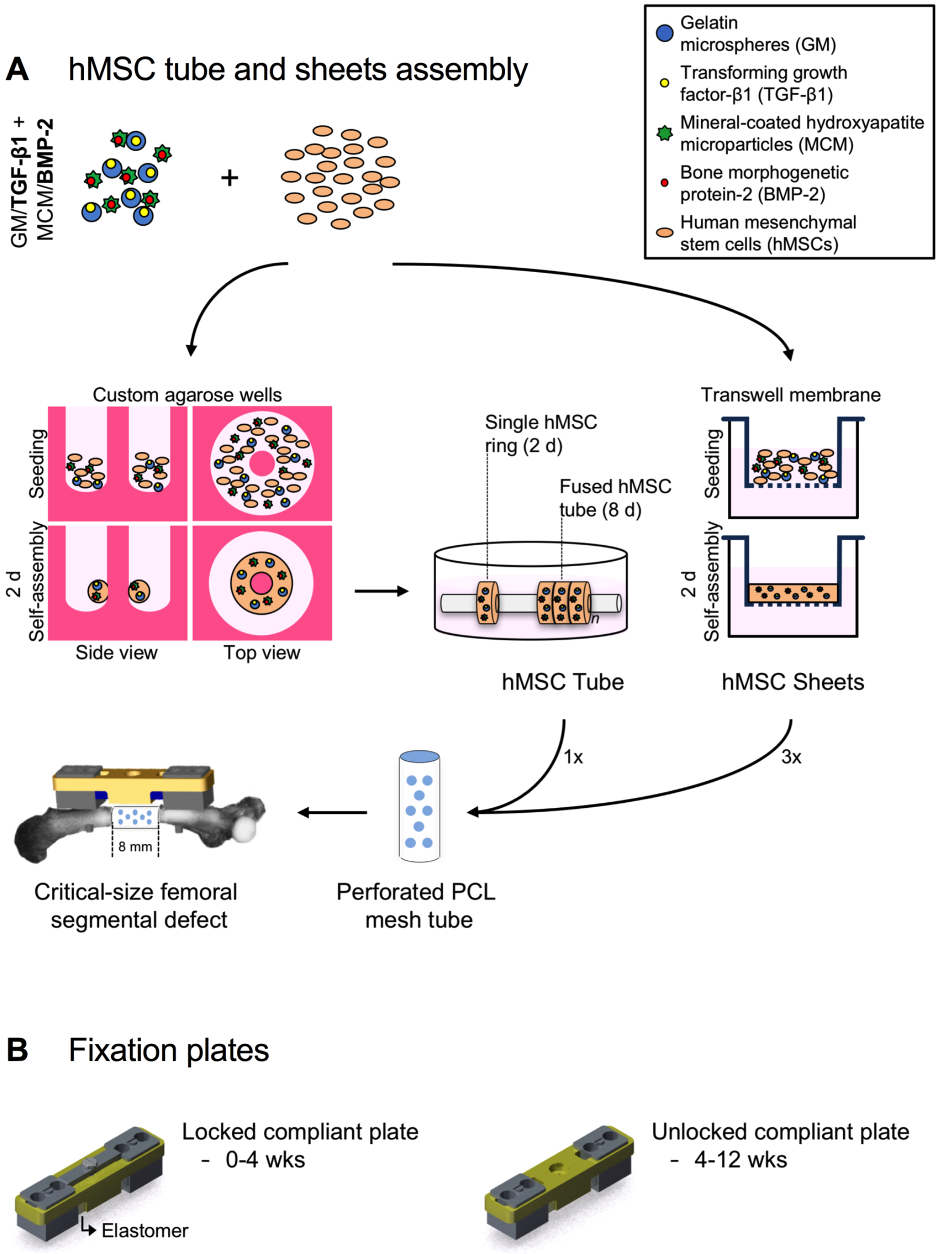


**Fig. S7. Schematic of hMSC condensate tube and sheet assembly for femoral defect implantation.** (A) hMSCs were mixed with TGF-β1-loaded gelatin microspheres and BMP-2-loaded mineral-coated hydroxyapatite microparticles [TGF-β1 + BMP-2], seeded in custom agarose culture wells and allowed to self-assemble into hMSC rings for 2 days before fusion into tubes by 8 days, or seeded onto membranes of transwell inserts and allowed to self-assemble into hMSC sheets for 2 days. One tube or three sheets (for identical cell number, microparticle concentration, and morphogen dose) were loaded into perforated polycaprolactone (PCL) nanofiber mesh tubes and implanted in critical-sized rat femoral segmental defects. (B) Limbs were stabilized with custom compliant fixation plates that were initially implanted in a locked configuration (0-4 weeks) to prevent load transfer, but were unlocked at week 4 to initiate ambulatory load transfer (4-12 weeks).

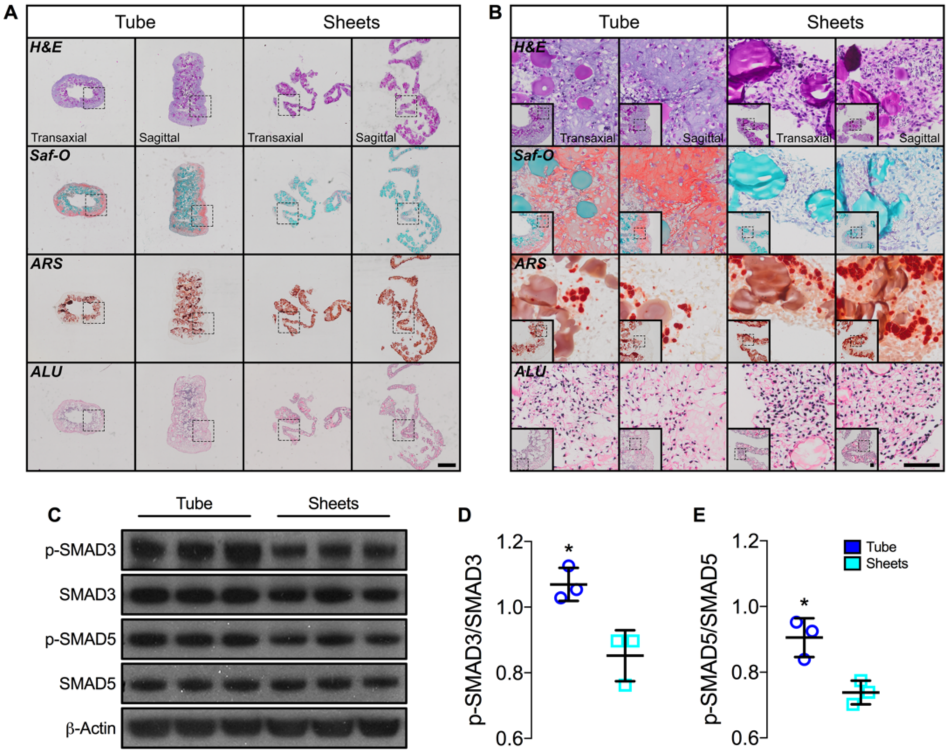


**Fig. S8. *In vitro* histological and biochemical evaluation of engineered hMSC condensate tube and sheet implants.** (A,B) Representative histological Hematoxylin & Eosin (H&E), Safranin-O/Fast green (Saf-O), and Alizarin Red S (ARS) staining, and *in situ* hybridization for human Alu repeats of transaxial and sagittal sections of hMSC tube and sheet implants containing TGF-β1 + BMP-2-loaded microparticles at day 8 and day 2, respectively. Scale bars, 1 mm ((A) dotted squares show areas used in 10x images in Fig. S8B) and 100 µm ((B) dotted squares in insets show region of interest in high magnification image). (C) Immunoblots and (D) relative quantification of p-SMAD3/SMAD3 and (E) p-SMAD5/SMAD5. β-Actin served as loading control (N = 3 per group; *$p<0.05$). Individual data points shown with mean ± SD. Analyzed by unpaired Student's *t*-test ($p<0.05$ or lower considered significant).

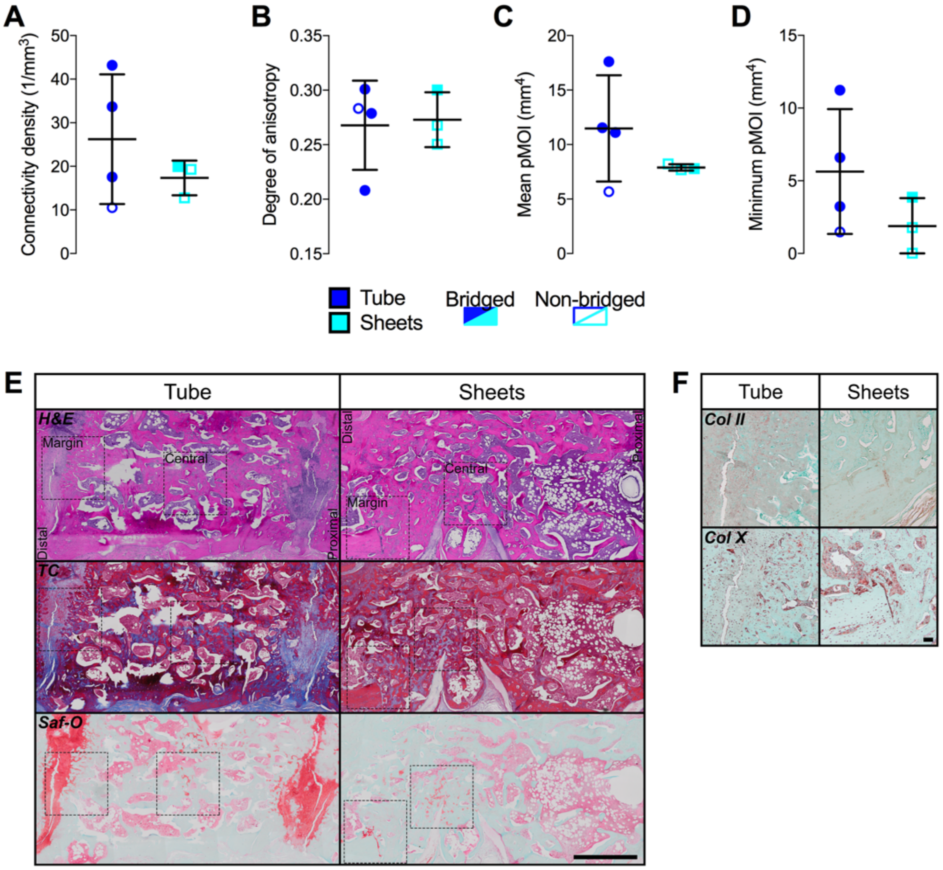


**Fig. S9.** ***Ex vivo*** **microCT, histological, and immunohistochemical evaluation of femoral defect healing induced by engineered hMSC condensate tubes and sheets.** Morphometric analysis of (A) connectivity density, (B) degree of anisotropy, (C) mean polar moment of inertia (pMOI), and (D) minimum pMOI in defects implanted with hMSC tubes or sheets containing TGF-β1 + BMP-2-loaded microparticles at week 12 (N = 3-4 per group). (E) Representative histological Hematoxylin & Eosin (H&E), Masson's Trichrome (TC), and Safranin-O/Fast green (Saf-O) staining of sagittal defect explant sections showing the complete defect; images oriented distal-to-proximal from left-to-right. (F) Representative immunohistochemical collagen (Col) II and Col X staining of sagittal hMSC defect explant sections showing the defect margin. Scale bars, 2 mm ((E) dotted squares show areas used in 10x images in Fig. 5G) and 100 µm ((F)). Individual data points shown with mean ± SD. Analyzed by unpaired Student's *t*-test ($p<0.05$ or lower considered significant).
